# Measuring single cell divisions in human cancers from multi-region sequencing data

**DOI:** 10.1101/560243

**Authors:** Benjamin Werner, Jack Case, Marc J. Williams, Kate Chkhaidze, Daniel Temko, Javier Fernandez-Mateos, George D. Cresswell, Daniel Nichol, William Cross, Inmaculada Spiteri, Weini Huang, Ian Tomlinson, Chris P. Barnes, Trevor A. Graham, Andrea Sottoriva

## Abstract

Cancer is driven by complex evolutionary dynamics involving billions of cells. Increasing effort has been dedicated to sequence single tumour cells, but obtaining robust measurements remains challenging. Here we show that multi-region sequencing of bulk tumour samples contains quantitative information on single-cell divisions that is accessible if combined with evolutionary theory. Using high-throughput data from 16 human cancers, we measured the *in vivo* per-cell point mutation rate (mean: 1.69×10^−8^ bp per cell division) and per-cell survival rate (mean: 0.57) in individual patient tumours from colon, lung and renal cancers. Per-cell mutation rates varied 50-fold between individuals, and per-cell survival rates were between nearly-homeostatic and almost perfect cell doublings, equating to tumour ages between 1 and 19 years. Furthermore, reanalysing a recent dataset of 89 whole-genome sequenced healthy haematopoietic stem cells, we find 1.14 mutations per genome per cell division and near perfect cell doublings (per-cell survival rate: 0.96) during early haematopoietic development. Our analysis measures *in vivo* the most fundamental properties of human cancer and healthy somatic evolution at single-cell resolution within single individuals.

## Introduction

Human cancers display tremendous inter-patient and intra-tumour genetic heterogeneity^1^. This heterogeneity is the consequence of a clonal evolutionary process marked by complex genomic changes^2^, parallel and convergent evolution^3^, non-cell autonomous dynamics^4^ and genomic instability^5^ that leads to metastatic spread, drug resistance and ultimately death^1,6,7^.

However, the microscopic forces underlying cancer evolution at the single cell level, such as the per-cell mutation rate and the per-cell survival rate remain immeasurable within individual human tumours^2,8^. Unlike species evolution for which a timed fossil record exists^9,10^, the lack of sequential data due to ethical and technical limitations is a major obstacle to quantitate somatic evolution in both healthy and cancerous human tissue. Moreover, high intra-tumour heterogeneity (ITH) necessitates measuring variation with extremely high precision, ideally at single cell resolution^3,4,11^. Precise single cell genomic measurements remain challenging and if possible can only be realized on a relative limited number of cells from tumours that may contain hundreds of billions of cells^12,13^.

Here we show that multi-region bulk samples of single tumours contain recoverable information about single cell divisions. Combining evolutionary theory, tumour multi-region sequencing and the ubiquitous stochastic nature of cell division and mutation accumulation can unravel this information.

### ITH in multi-region data encodes the properties of single-cell divisions

All tumour cells from a tumour bulk sample descended from a most recent common ancestor (MRCA) cell. In a tumour, typically composed of hundreds of billions of cells, multiple spatially separated bulk samples differ in their exact composition of somatic mutations, because mutations accumulate across different branched cell lineages during growth ^14,15^, (Figure 1A and Methods). Branching is inevitable in evolutionary processes driven by cell division and mutation, both in the presence and absence of clonal selection^1,6,7,15–17^.

**Figure 1:**
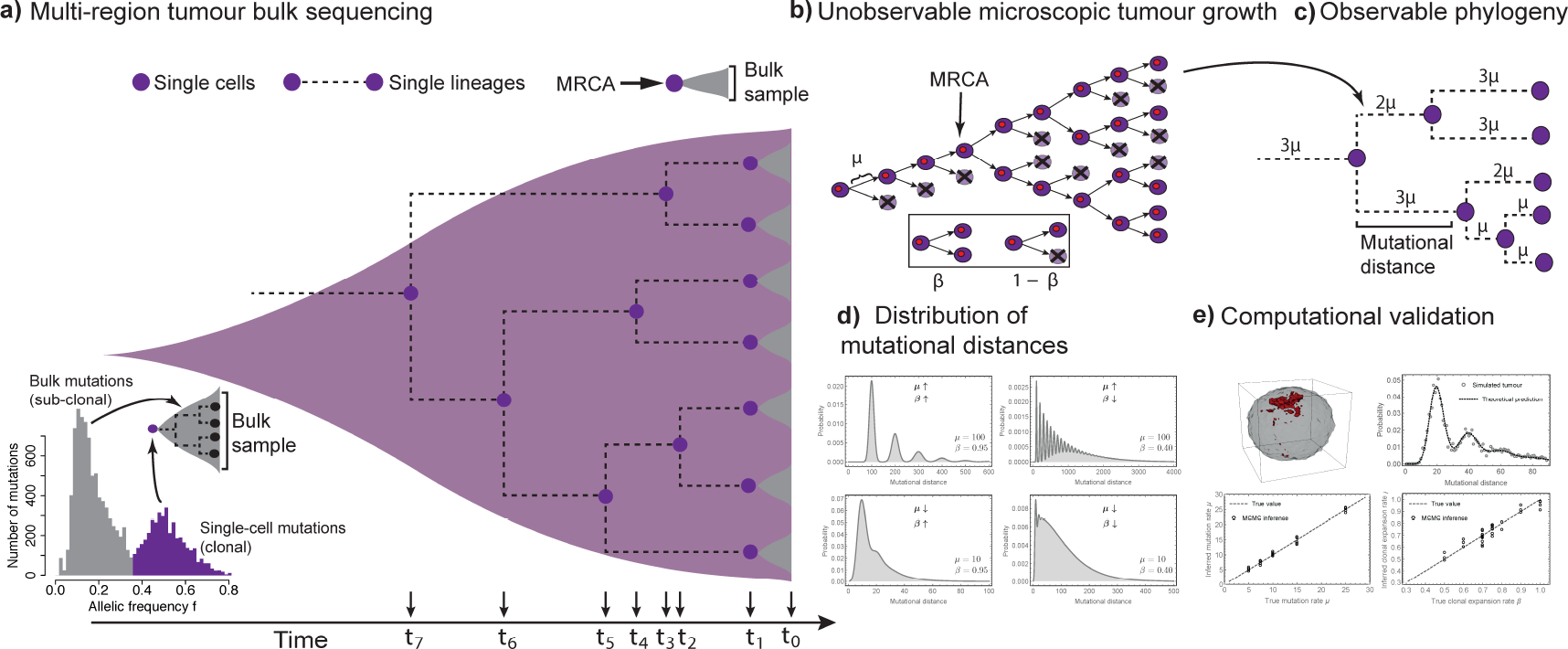
Multi-region tumour bulk sequencing encodes information on single cell lineages and single cell divisions. **a)** Each of the seven spatially separated tumour bulk samples (in grey) consists of thousands to millions of cancer cells that descended from a single most recent common ancestor (MRCA) cell. The genomic make-up of the single ancestral cell is described by the mutations clonal to the bulk sample. Those appear at high variant allele frequency in the sample (bottom-left panel, in purple). The intersection of mutations in any two bulk MRCA cells corresponds to the genomic profile of another more ancestral cell. This process continues back in time until the MRCA cell of all the sampled cells is reached. **b)** The level of genomic variation within a tumour is the direct consequence of mutation accumulation during cell divisions, leading to complex branching structures. Intervening selective pressures, trimming certain branches while favouring others, may further modify these structures. Importantly, the most fundamental parameters, the per-cell mutation rates and per-cell survival rate that drive this process are not directly observable. **c)** Per-cell mutation rate per division *μ* and per-cell survival rate *β* leave identifiable fingerprints in the observable patterns of intra-tumour genomic heterogeneity. Cell divisions occur in increments of natural numbers and thus the mutational distance between any two ancestral cells is a multiple of the mutation rate *μ*. **d)** The quantized nature of cell divisions leads to a characteristic distribution of mutational distances across cell lineages. The shape of the distribution depends on the exact values of *μ* and *β*. Roughly four different scenarios of small and large *μ* and *β* are possible. Importantly, they influence the shape of the distribution differently and thus constructing the distribution of mutational distances allows disentangling the per-cell mutation rate *μ* and per-cell survival rate *β*. **e)** Spatial stochastic simulations of growing tumours confirm the ability of mutational distance distributions to disentangle mutation and lineage expansion rates. A Monte Carlo Markov Chain framework based on mutational distance distributions reliably identifies mutation and lineage expansion rates in simulations of spatial and stochastically growing tumours (*μ*: Spearman Rho = 0.98, *p* = 4×10^−23^; *β*: Spearman Rho = 0.93, *p* = 8×10^−16^, Relative error: *η*_*μ*_ = 0.056, *η*_*β*_ = 0.045).

The *mutational distance*, the number of somatic mutations different between two ancestral cells, emerges from two dynamic processes: (i) the per-cell intrinsic mutation rate per division *μ*, and (ii) the number of cell divisions separating ancestral cells in space and time. The latter depends on the per-cell survival rate *β*, or in other words the probability for a single cell division to establish two surviving lineages. Hence, the per-cell survival rate *β* accounts for lineage loss due to cell death or differentiation (Figure 1b,c). *A priori*, both *μ* and *β* are unknown. Previous methods measure effective mutation rates 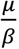, but cannot entangle these for evolution fundamental microscopic parameters ^15,17–19^.

However, cell divisions must occur in increments of natural numbers (cells are 1, 2, 3,…,n cell divisions apart), whereas the cell intrinsic mutation rate follows a Poisson distribution (Figure 1d). As described in the Methods, the distribution of mutational distances in a tumour encodes these two properties of single cell divisions. This is possible because many ancestral cells are only a few cell divisions apart (SI Figure 19). Specifically, we show that the probability density of mutational distances *y* in a tumour takes the following form:

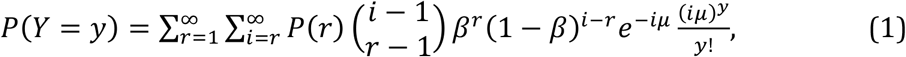

given a per-cell mutation rate *μ* and a per cell survival rate *β* (Methods). Equation (1) predicts four possible regimes for the distribution of mutational distances, discriminated by uni- or multimodality determined by the combination of small or large *μ* and *β* (Figure 1d). The parameters uncouple in above equation and thus repeated sampling from the distribution allows measuring both parameters separately. Importantly, this can only be done when enough (≥ 6) bulk samples are available for each tumour (Figure 1d & SI Figure 33). Moreover, our approach relies on comparing mutational distances between samples and does not require *a priori* clonal decompositions of tumours. We demonstrate that individual based stochastic simulations of spatial tumour growth converge to the abovementioned analytical solution (Figure 1e & Methods) and a Bayesian inference scheme recovers the imposed evolutionary parameters (Figure 1e and Methods). We also demonstrate that our approach remains robust when the underlying assumptions are relaxed, e.g. non-constant mutation rates or the presence of subclonal selection during population expansion (see Methods). Moreover, we also analyse how the sensitivity of the evolutionary estimates depend on the quality and quantity of the genomic data (see Methods).

**Figure 2:**
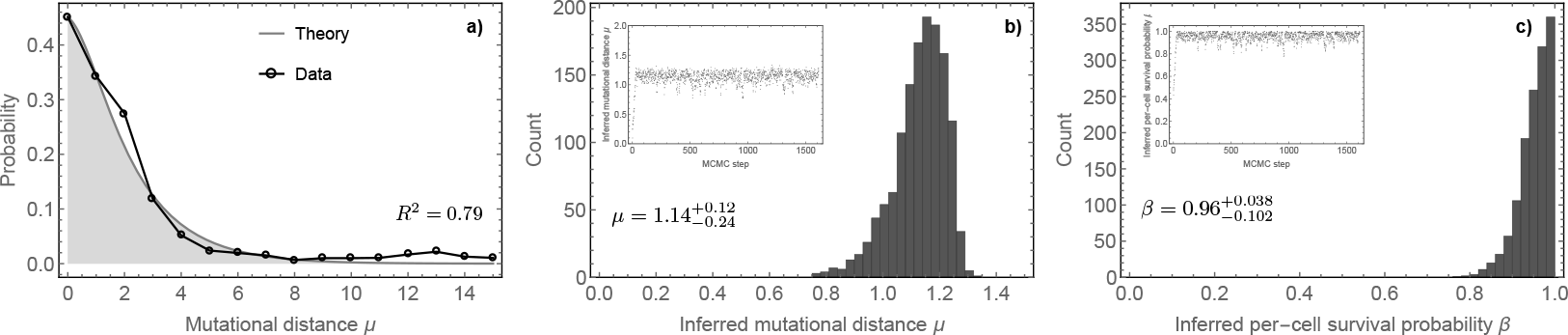
Per-cell mutation and per-cell survival rate inferences in healthy haematopoiesis during development. **a)** Mutational distance distribution inferred from 89 whole genome sequenced healthy haematopoietic stem cells (black dots), data taken from^20^ and best theoretical fit (grey line). MCMC inference for **b)** the mutation rate per cell division (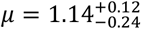 mutations per whole genome per cell division) and **c)** the per-cell survival rate 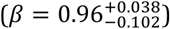 during early development in healthy haematopoiesis. Median values and 95% credibility intervals were taken from the posterior parameter distributions.

### Per-cell mutation and per-cell survival rate in healthy haematopoietic development

Before we discuss properties of individual tumours, we tested our approach in a biologically well-characterised *in vivo* example of somatic evolution. We make use of the accumulation of somatic mutations during early development of the healthy haematopoietic system as a benchmark. In a recent article Lee-Six and colleagues whole genome sequenced 89 healthy haematopoietic stem cells of a single 59 year old male ^20^. They subsequently constructed the phylogeny of healthy haematopoiesis and estimated the per-cell mutation rate to be 1.2 mutations per genome per division during early development assuming perfect cell doublings. We can use the same sequencing information and construct the distribution of mutational distances during early haematopoietic development. Our framework of mutational distances allows a joint and independent inference of the per-cell mutation and per-cell survival rate (Figure 2 and Methods). In agreement with ^20^, we find a median mutation rate of 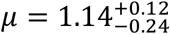 mutations per genome per division (shown is the medium mutation rate per bp/cell-division and 95% credibility intervals). Furthermore, we infer a per-cell survival rate of 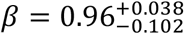, independently confirming the original assumption of almost perfect cell doubling during early development ^20^.

### Measuring the per-cell mutation rate in individual human tumours

We now proceed to *in vivo* per-cell mutation rates within individual human tumours. A unique sample set^21^ amenable to our analysis is composed of 6 colorectal tumours (5 carcinomas, 1 adenoma) sequenced using multi-region whole genome (3 tumours) and whole exome profiling (3 tumours) of up to 13 bulk samples per tumour (median 7.6, minimum 6 samples per tumour as required by our analysis). We calculated the pairwise genetic divergence for all combinations of samples per tumour and used our MCMC approach to infer the per-cell mutation rate *μ* as well as the per-cell survival rate *β* from equation (1) above. Simulations show that inferences are possible with as few as 6 tumour samples (SI Figure 33). Despite the limited resolution (median 8 bulk samples per tumour), our theoretically predicted mutational distance distribution describes important features of the data well (Figure 3 and SI Figure 13). Similar distributions emerge in stochastic simulations of tumour growth with comparable data quality (SI Figure 34). When whole genome sequencing was available, the mutational load was sufficient to apply the inference framework to each chromosome separately (Figure 3 and SI Figures 1–3 & 9–18). The analysis was restricted to regions of chromosomes with same copy number profile in all samples of a tumour and inferences were normalised by copy-number and genome content sequenced.

We found mutation rates per cell division to be elevated approximately 10 to 30 times compared to healthy somatic tissue^10^ across the whole genomes of the three carcinomas 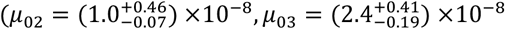 and 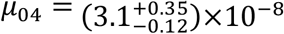 bp/division, median mutation rate and 95% credibility intervals), see Figure 3 and Methods. Mutation rates differed significantly between tumours, but not across chromosomes of the same tumour (SI Figure 15). Recently it was suggested that mismatch repair efficacy differs between coding and non-coding genomic regions^22^. We find mutation rates in coding compared to non-coding regions slightly elevated in mismatch repair sufficient tumour 02 and slightly lower in tumour 03, but being equal in the mismatch repair deficient tumour 04 (SI Figure 5). We found comparably high mutation rates per cell division for exonic mutations in two additional carcinomas (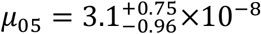 bp/division and 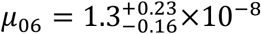 bp/division), SI Figure 16. Instead, one adenoma showed a near normal per-cell mutation rate (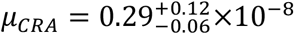 bp/divison), SI Figure 16. Overall this suggests important differences in mutation accumulation at the single cell level between tumours and is in good agreement with recent experimental *in vitro* single cell mutation rate inferences^23^.

**Figure 3:**
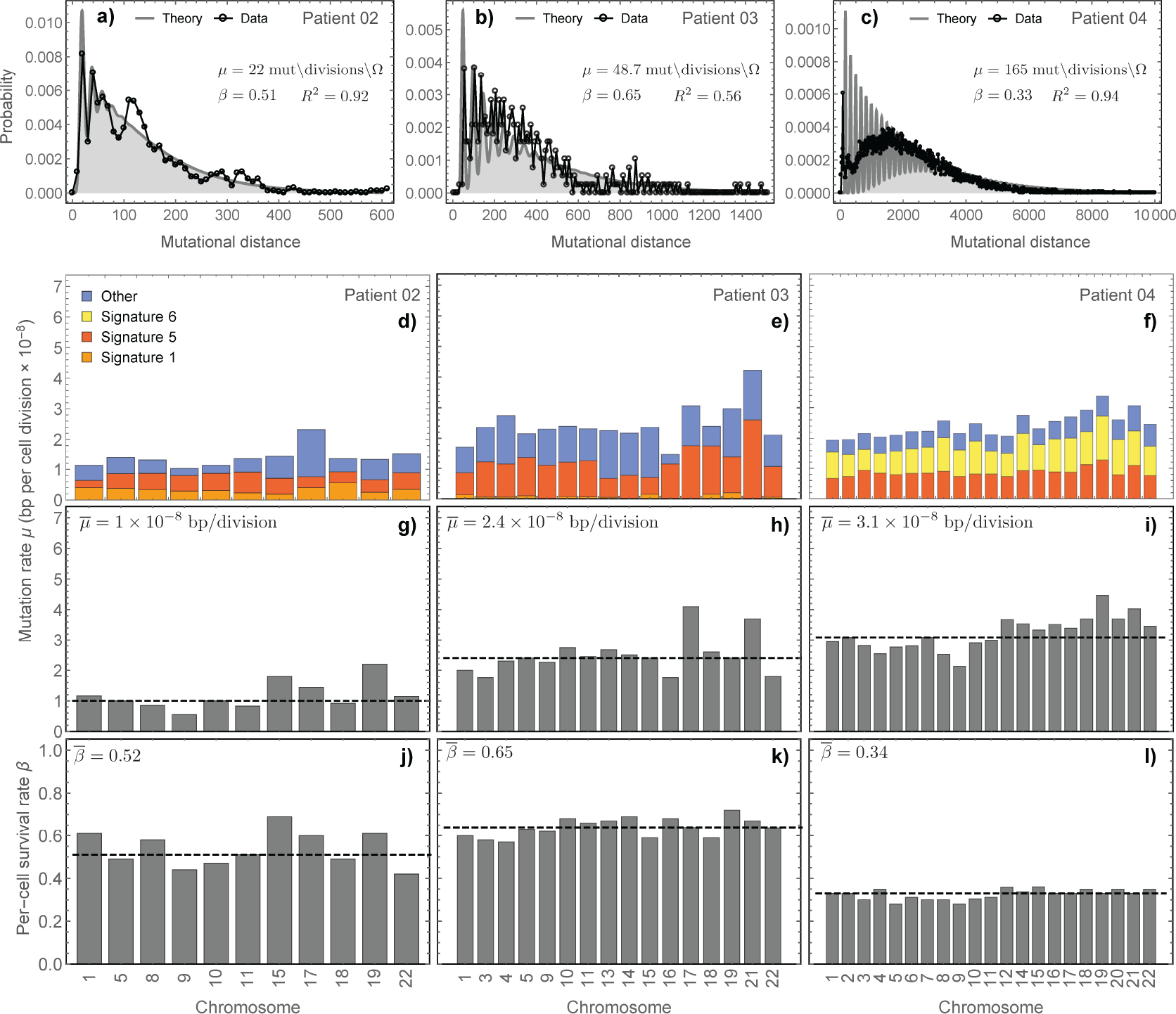
Mutational distance distributions reveal per-cell mutation and per-cell survival rates. **a-c)** Mutational distance distributions (whole genome) for three colorectal carcinomas21 (dots=data, grey line=theoretical prediction based on MCMC parameter estimates). Patient 04 (MSI+) has one order of magnitude larger mutational distances. Three additional distributions are shown in SI Figure 13. **d-f)** Per-cell signature mutation rate per chromosome. Results are consistent across chromosomes (Methods). **g-i)** The median overall mutation rates are 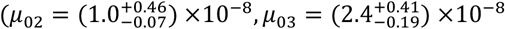 and 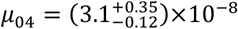 bp/division, dashed lines), 10 to 30 times higher compared to healthy somatic cells. Patient 04 is MSI+ highlighted by signature 6. **j-l)** Estimates of per-cell survival rates per chromosome are consistent across chromosomes of the same patient (Median: 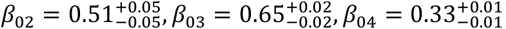), but vary considerably between patients (SI Figure 15).

To further unravel the underlying differences in mutation accumulation, we decomposed somatic mutations into the most prevalent mutational signatures^24^ for all three whole-genome sequenced colorectal carcinomas and inferred per-cell mutation and per-cell survival rates per signature in each chromosome (Figure 3 and Methods). Signature 5 was detected consistently across all chromosomes for all three carcinomas, but the accumulation rate differed between tumours (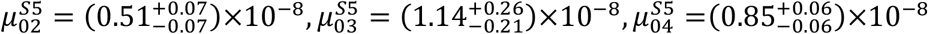 bp/division). Signature 1 was identified in all chromosomes of tumours 02 and 03 and was similar to a healthy somatic mutation rate (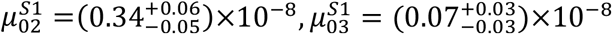 bp/division), further supporting its previously proposed clocklike nature in aging human tissues^25^. Consistent with its classification as MSI+, Signature 6 was prominent in tumour 04 (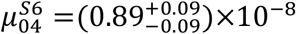 bp/division), comprising 39% of all mutations in the tumour (63% if mutations of unclassified signatures are included). All somatic mutations not assigned to abovementioned signatures were grouped as other (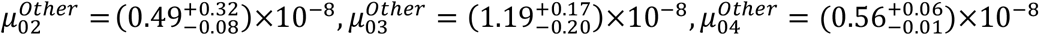 bp/division). Overall our analysis suggests variation in between patient signature mutation rates (SI Figure 7). In contrast, we do not find significant dependence of per-cell mutation rates on chromosomal ploidy: mutation rates remained consistent for diploid, triploid and tetraploid chromosomal regions after correcting for the genome size sequenced (SI Figure 8).

The differences in mutational signatures between individuals also manifest in variable accumulation rates of substitution subtypes (SI Figure 6). In two individuals, transitions are more likely than transversions. As expected C → T transitions have the highest mutation rates, but accumulation rates differ between individuals. Interestingly, the ranking of the rates of substitution subtypes in two patients agree with the patterns of divergence between humans and chimpanzees^26^. In contrast, in one patient mutation rates for transitions and transversions were similar (SI Figure 6b).

In addition, we inferred per-cell mutation rates per chromosome in 3 recently published^27^ whole genome sequenced colon cancer patients (7, 9 and 9 tumour samples). We also used data from two non-small cell lung cancers (NSCLC) from the TRACERx study^28^. These were the two cases (one squamous and one adenocarcinoma) that had more than 6 samples per tumour from the 100 patients cohort (7 exome sequenced samples each), as well as five clear cell renal cell carcinomas (CCRCC)^29^ (median 8, from 8 to 12 exome sequenced bulk samples), (SI Figures 9–12 & 17,18). In concordance with our previous observation we found consistent mutation rates across chromosomes for the colon cancer patients. One MSI+ case has an increased mutation rate (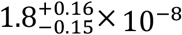 bp/divison) compared to two MSS patients (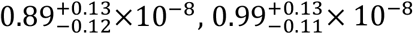 bp/divison). We found the lung squamous cell carcinoma to have a very high mutation rate (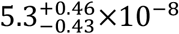 bp/division), in comparison the lung adenocarcinoma had a lower mutation rate (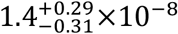 bp/division). Also, three clear cell renal carcinomas showed elevated mutation rates (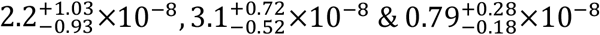 bp/division). Surprisingly, two clear cell renal carcinomas had near normal somatic mutation rates (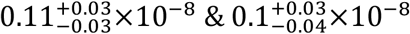 bp/division), suggesting that at least in some cases, cancer cells maintain near normal mutation rates per cell division.

### Measuring the per-cell survival rate in individual human tumours

Our inference scheme allows a joint estimate of the mutation rate per cell division and the per-cell survival rate. We observed striking differences for the per-cell survival rates between the tumours discussed above. We found for the colorectal tumours (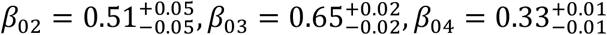), where higher *β* corresponds to less cell death. The rates were consistent when the analysis was based only on individual chromosomes (Figure 3). Interestingly, tumour 04 was mismatch repair deficient and had 3 to 10 times higher sub-clonal mutational burden compared to mismatch efficient carcinomas (SI Figure 4), but remarkably slower growth. Hence, the higher mutational load in this tumour may not solely be due to mismatch repair deficiency, but also due to slower growth and therefore older relative tumour age (more cell divisions). This is also consistent with clinical observations and may explain partially why MSI+ tumours are more prevalent in older patients and typically have a better prognosis^30–32^. Per-cell survival rates inferred from exonic mutations also varied for two additional carcinomas 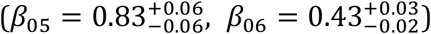 and was at the lower bound for the adenoma 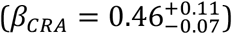. The 3 independently whole genome sequenced colon cancers^27^ show similar per-cell survival rates 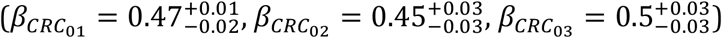. The lung squamous cell carcinoma had a very low per-cell survival rate 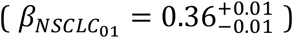. Interestingly, the lung squamous cell carcinoma and the two MSI+ colorectal cancers have amongst the highest mutation rates but the lowest per-cell survival probabilities (Figure 4a). In comparison, the lung adenocarcinoma had a higher per-cell survival rate 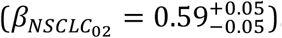. All but one clear cell renal cell carcinoma had high per-cell survival rates 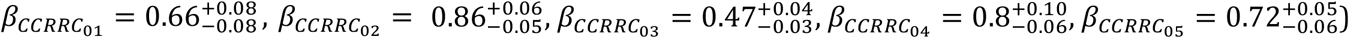.

**Figure 4:**
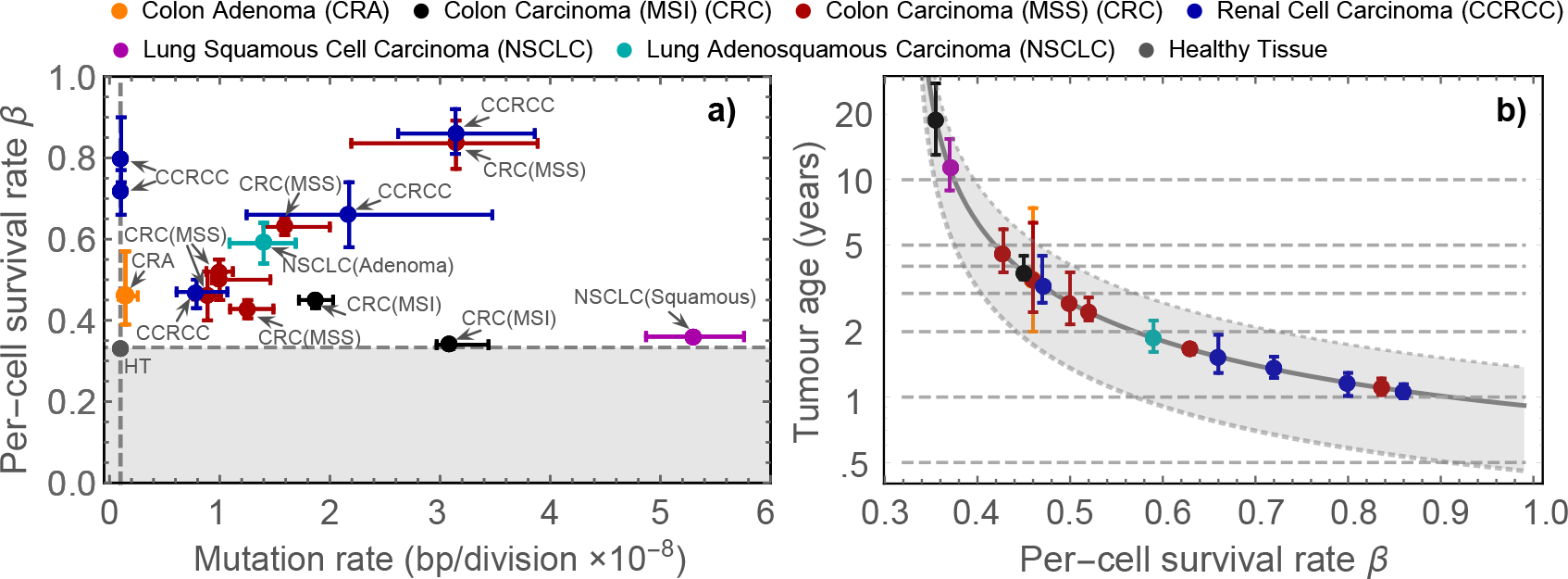
Map of per-cell mutation and per-cell survival rates across cancer types. **a)** The intersection of the dashed lines correspond to values of healthy tissue during homeostasis (*μ*_*h*_ = 1×10^−9^, *β*_*h*_ = 1/3). White background corresponds to values of *β* that allow for growing cell populations, shaded area describes values of *β* that would lead to population extinction (see Methods). Error bars show 95% credibility intervals. There are three different patterns, a subset where lineage expansion and mutation rates correlate positively (SpearmanRho =0.85, p=0.002, excluding 6 samples with near normal mutation or near normal survival rate), a subset of cases with near normal mutation and another group with near normal per-cell survival rates. **b)** The per-cell survival rate *β* can be translated into tumour age at diagnosis (duration from tumour initiating cell to diagnosis). We find that most tumours are 1 to 5 years old (Median: MSS Colorectal Carcinomas: 2.6 ±1.5 years, Renal cancers: 1.34 ± 0.9 years, Colorectal Adenoma: 3.5 years, MSI+ Carcinoma: 18.8 and 3.8 years, Lung adenocarcinoma: 1.9 years, Lung squamous cell carcinoma: 11.3 years). Error bars show 95% credibility intervals, the grey line assumes a lineage division rate once every 2 weeks, the grey area corresponds to division rates of once every 1 to 3 weeks respectively.

Figure 4a shows each tumour’s per-cell survival rate *β* plotted against its per-cell mutation rate *μ*. Healthy homeostasis implies an overall constant cell population, corresponding to *β* = 1/3, and an approximate somatic mutation rate of *μ* = 1×10^−9^ (bp/division) (Figure 4a). Tumours distribute widely across these evolutionary measures, emphasizing the uniqueness of each individual tumour. The adenoma is overall most similar to healthy somatic tissue. Interestingly, there seem to be 3 distinct scenarios. In some tumours, per-cell survival and mutation rate are positively correlated (SpearmanRho =0.85, p=0.002, excluding 6 samples with near normal mutation or near normal survival rate). However, there is a subset (3 out of 16) of tumours with accelerated growth and near normal somatic mutation rates and another group (3 out of 16) with high somatic mutation rates but near normal per-cell survival rates (Figure 4a).

### Estimating tumour age at diagnosis

The per-cell survival rate inferences allow approximations for the duration of tumour expansions across patients and cancer types. Assuming 10^11^ tumour cells at diagnosis, tumour ages are between 30 and 600 generations for the fastest growing chromosomally unstable carcinoma and the slowest growing MSI+ carcinoma. For lineage division rates of once every two weeks, the duration of the final expansions are between 1 and 19 years (Median: MSS Carcinoma: 2.6 ±1.5 years, Renal cell carcinoma: 1.34 ± 0.9 years, Adenoma: 3.5 years, MSI+ Carcinoma: 18.8 and 3.8 years, Lung adenocarcinoma: 1.9 years, Lung squamous cell carcinoma: 11.3 years), see Figure 4b. These estimates correspond to the duration of the final phase of cancer cell expansion and remain within a 10% error bound for a one order of magnitude deviation of the tumour size at diagnosis (SI Figure 23). Ranges for different lineage division rates are shown in Figure 4b.

## Discussion

Here we have shown how the mutational burden, now routinely measurable in healthy and cancerous tissues, emerges from intertwined microscopic evolutionary forces, the per-cell mutation and per-cell survival rate. More importantly, multi-region tumour sequencing allows a joint inference of these forces and reveals major differences between individual patients. Furthermore, evolutionary forces may intertwine with other cell intrinsic and extrinsic processes and may or may not change in time. Unravelling these interactions will require further more fine-grained sampling of tumours, ideally both in time and space. Sequencing of potentially thousands of single cells promises a significant information gain that will allow for much higher resolved mutational distance distributions in the near future. Nevertheless, it seems that inferences of tumour evolution and subsequent treatment strategies solely based on population averages risk error prone conclusions for any individual patient. A personalised unravelling of the microscopic evolutionary forces appears essential.

## Methods

### The distribution of mutational distances

Multi-region bulk sequencing of tumours allows us to reconstruct the evolutionary history of single cell lineages, see also Figure 1 in the main text. Each tumour bulk sequence contains information about clonal and sub-clonal mutations. Clonal mutations of a bulk sample are present in all cells of the sample and therefore must have been originated from a single joined most recent common ancestor cell that gave rise to all sampled cells in the tumour bulk. In contrast, sub-clonal mutations are present in a subset of cells in the tumour bulk and arose later during tumour growth. Consequently, multiple bulk sequences allow reconstructing the genomic composition of multiple single cells that existed at different times during the life history of the growing tumour population. This principle allows us to apply phylogenetic methods to cancer genomic data. Mutations that distinguish most recent common ancestor cells were accumulated during a finite number of cell divisions (cell divisions are necessarily quantised). During each cell division, daughter cells might acquire additional novel mutations. The number of novel mutations X after a single cell division depends on the mutation rate *μ* and the length of the genome *L*. The number of novel mutations per cell division *X* follows a *Poisson* distribution

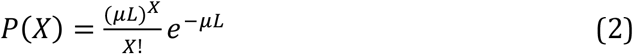

with mean and variance *μL*. Usually, the length of the sequenced genome *L* is known (for example the exome or whole genome of a cancer cell) and the mutation rate per cell division *μ* is the only unknown parameter. Thus sampling sufficiently many mutational distances of single cell divisions allows us (in principal) to reconstruct the underlying *Poisson* distribution and therefore the inference of the mean mutation rate per cell division *μ*. However, distances between cells of a lineage might be larger than a single cell division and double, triple and higher modes of cell division contribute to the distribution of mutational distances of multi-region samples. For example, if a cell divides twice, it will acquire novel mutations twice and the total number of mutations *X*_1_ + *X*_2_ is the sum of two independently *Poisson* distributed events *X*_1_ and *X*_2_. The number of novel mutations *X*_1_ + *X*_2_ is again *Poisson* distributed, but now with mean 2*μL*. In general, a cell accumulates *X*_1_ + *X*_2_ + … + *X*_*n*_ *Poisson* distributed number of novel mutations

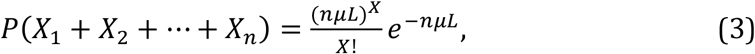

after *n* cell divisions. However, we must also account for cell death or differentiation, leading to lineage extinction. Indeed, branching in the evolutionary history of the tumour only occurs if both daughter cells survive. We therefore introduce a probability *β* of having two surviving lineages after a cell division and a probability 1 − *β* of a single surviving lineage respectively. Thus, *r* cell divisions with two surviving lineages (successful divisions) are accompanied by *m* cell divisions with only a single surviving lineage (unsuccessful divisions).

The number of unsuccessful divisions *m* can be understood as a random variable again. More specifically, they follow a Negative Binomial distribution

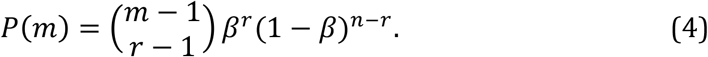

Thus the number of mutations acquired between two successful divisions depends on the *Poisson* distributed mutation rate *μ* and the Negative binomial distributed number of unsuccessful divisions *m*. Intuitively, a certain measured mutational burden in a single cell lineage or bulk sample of a tumour can result either from many unsuccessful divisions with a low mutation rate or, alternatively a few unsuccessful divisions with high mutation rate. Formally, we can write for the total number of mutations between two successful divisions

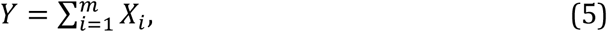

where *X*_*i*_ are independently distributed *Poisson* random variables and *m* is a Negative binomial distributed random variable.

Now we can seek the probability of the number of acquired mutations *Y*_*r*_ after *r* successful divisions. We first note that the probability generating functions for both *Poisson* and Negative Binomial distributed random variables are known and given by

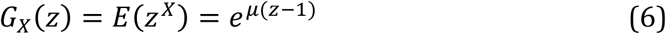

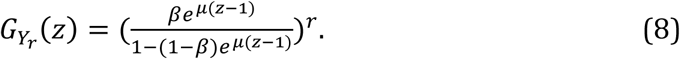

Using these expressions and the law of total probability, this implies for the probability density function of the joint distribution *Y*_*r*_

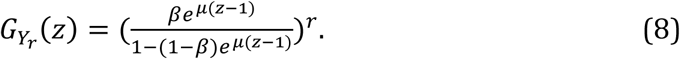

Finally, we are interested in the probability *P*(*Y*_*r*_ = *y*), for example the probability to observe a certain mutational load *y* given a mutation rate *μ*, a number of successful divisions *r* and a survival rate *β*. We can expand the probability generating function into a power series and write

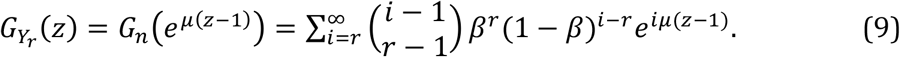

Expanding the exponential function, we can write

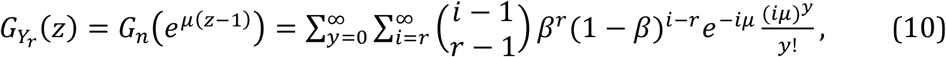

and thus, we find for the probability of having *y* mutations after *r* successful divisions

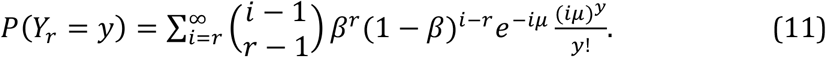

A complete description of the distribution of mutational distances requires an expression for the expected distribution of successful divisions *r* (the number of branching events between two cell lineages). Remaining general, we can write

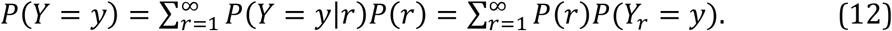

Substituting equation (11) the probability density for the mutational distances becomes

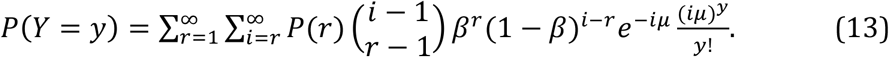

Note, the expected distribution of successful divisions *P*(*r*) is independent of the underlying mutation rate *μ*, it only depends on the per-cell survival probability *β*. It therefore does not impede our ability to disentangle *μ* and *β*. In the following we derive an explicit expression for *P*(*r*) for an exponentially growing population.

### The distribution of successful divisions *r* for an exponentially expanding population

Expanding on classical results of coalescence theory^33^ we can derive an analytical expression for the distribution of successful divisions *r* in the case of an exponentially growing cancer cell population. Assume a population of cancer cells grows exponentially in time with *N* (*t*) = *N*_0_*e*^*λβt*^. Here *β* corresponds to the survival probability of two lineages that was introduced above and time *t* is measured in generations. We are interested in events backward in time *t* → −*t* and thus our population effectively shrinks exponentially *N* (−*t*) = *N*_0_*e*^*−βt*^. The probability of coalescence of two cells at time (*t*) given no coalescence before *t* is approximately 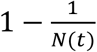 and the probability to coalesce at time *t* is 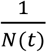 Thus the probability that the first coalescence occurs at exactly time *t* is approximately given by

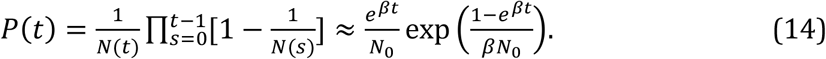

In our case, we are concerned with mutational distances and thus we ask for the distribution of times between coalescence events Δ*t* rather than the distribution of coalescence time *t*. However, we can directly infer this distribution from equation (14), by rewriting Δ*t* = *t*_0_ − *t* as the time of the initiating cell population at some point in the past. By substituting *t* = log *N*_0_ /(*β*) we have 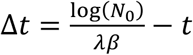 and we find for the distribution of times between coalescence events

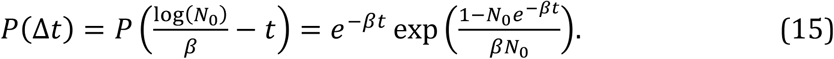

This is for sufficiently large *N*_0_ well approximated by

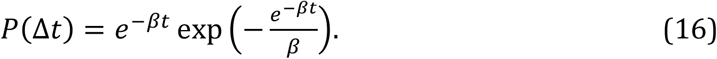

Examples of Equation (15) and (16) are shown in SI Figure 20. We can discretise this probability density function to derive at the probability for the number of successful divisions *r* via

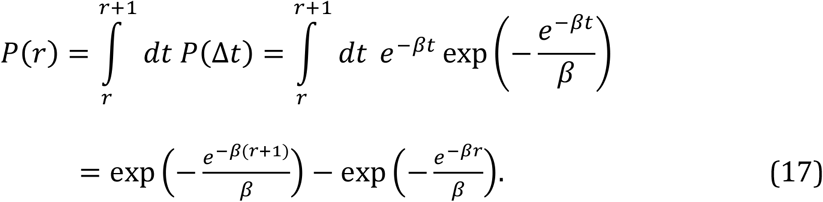

As we are interested in positive branch length only, we need to normalise the distribution for non-negative integers. The normalising factor is 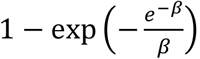 and the distribution of successful divisions *r* in an exponentially expanding cell population becomes

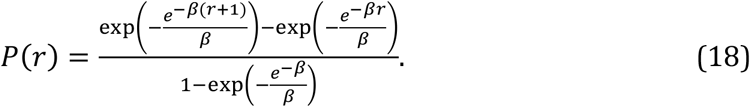

In combination with equation (13) this allows a complete description of the distribution of mutational distances in exponentially growing cell populations, SI Figure 21. Interestingly, equation (18) predicts that on average most successful divisions are only a few cell divisions a part (SI Figure 19).

This probability density fits the distribution of mutational distances from simulated tumour growth exactly (Figure 1e). Furthermore, utilising MCMC methods, this distribution allows solving the inverse problem. We can infer the single cell parameters (mutation rate *μ* and cell survival rate *β*) from one time measures of the distribution of distances of multiple samples from a single tumour.

### Properties of the mutational distance distribution

We introduced the distribution of mutational distances as a measure to disentangle per-cell mutation and per-cell survival rates. This distribution has 2 free parameters that determine its shape (*μ*: the number of mutations per cell division and *β*: the per-cell survival probability). The combination of these 2 parameters allows for different shapes of the distribution and predicts 4 possible distinct scenarios given by the combination of small and large *μ* and *β*. Examples for the theoretically expected shape of the mutational distance distribution are shown in SI Figure 21. A multi-modal distribution is expected for sufficiently large mutation rates per cell division *μ* (SI Figure 21a & b). In contrast, uni-modal distributions become evident for smaller mutation rates *μ* (SI Figure 21c & d). The per-cell survival probability *β* determines the height of the modes as well as the length of the tail (emphasized by at least one order of magnitude differences in the y- and x-axes of the panels in SI Figure 21). In SI Figure 34 we show that these scenarios are reproduced in stochastic spatial simulations of tumour growth. Also note, in SI Figure 34 we plot examples of the mutational distance distribution (with fixed *μ* and different *β*) at scale to emphasize some of the significant differences.

### Spatial intermixing, sequencing noise and the number of mutational distances

Our method of mutational distances does not require the construction of phylogenetic trees. Instead, it is based on pairwise mutational distances (the number of mutations that separate ancestral cells) and thus we construct pairwise differences of mutational counts.

The mutational load of ancestral cells is constructed from intersections of the complete lists of mutations of all combinations of bulk samples. The mutational distances correspond to the number of unique mutations between any two such intersections.

If we were sampling on a tree of perfectly defined species, the number of inferable intermitted branches would be in the order 2*n* − 2, given *n* species. However, our situation is slightly different. A spatial bulk cancer sample contains multiple lineages of cells that are sequenced. Due to cell intermixing and subclonal mutations, groups of cells in the same sample may have distinct common ancestors. In SI Figure 22 a we show that, if we were able to sequence at perfect clonal single cell resolution, we would indeed infer 2*n* − 2 uniquely different intermittent mutational distances. However, allowing realistic spatially sampled bulk populations with lineage intermixing results in more identifiable ancestral populations and thus more mutational distances (SI Figure 22). Given *n* independent bulk samples, we can split the data into smaller sets containing *i* samples. For each such set we have 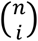 possible intersections and the possible number of intersections between *i* − 1 and *i* subsets becomes 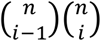. Consequently, the maximal combinatorial number of mutational distances given *n* bulk samples is

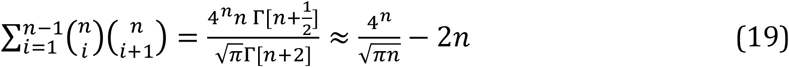

and scales faster (leading term ~4^*n*^) compared to the number of intermittent branches of a binary time ordered tree (~2*n*) (SI Figure 23). This is also exemplified in SI Figure 22, which compares the mutational distance distribution given a perfectly inferable clonal structure (species tree) to the case of spatial tumour bulk sampling with lineage intermixing and noise from sequencing depth. It has to be said that not each of these distances is unique and we will sample some of the same distances repeatedly. However, the extractable information is sufficient to recover the theoretically predicted distribution of mutational distances (Figure 1e and SI Figures 22, 27, 28 & 34).

### Connection of the effective survival rate and a microscopic death rate

Throughout our derivation of the mutational distance distribution we use an effective survival rate *β* to model cell death. We defined *β* as the probability that both cell lineages survive after cell division, while 1 − *β* is the probability of only a single surviving cell lineage. This concept is closely related to cell fitness, as cells with higher *β* have on average more successful cell divisons and thus produce (given the same number of cell divisions per unit time) more surviving offspring compared to cells with lower *β*. On could also formulate cell death with a microscopic perspective that would suggest a certain probability of cell death *α* for each daughter cell after division. Such a probability would allow three outcomes after a cell division: with probability (1 − *α*)^2^ both daughter cells survive, with probability 2*α*(1 − *α*) one daughter cell survives and with probability *α*^2^ both daughter cells die. However, as we are bound to find surviving cell lineages in every possible measure of tumours, none of the observed cell lineages can have gone extinct. Thus an event where both daughter cells died (cell lineage extinction) did not occur within the observable data. Mathematically, this implies that every measurement conditions cell division on non-extinction of both daughter cells and we can write

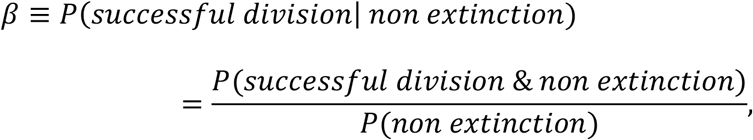

and with the corresponding probabilities *α* we get

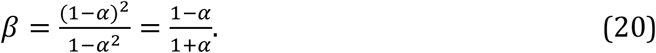

In our scenario *β* and *α* are interchangeable. We also can rearrange equation to solve for *α*,

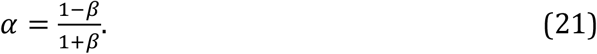

If we interpret *α* as the probability of random cell death after a division, *α* must be smaller than 1/2. If *α* were larger than 1/2, tumour populations extinct almost surely after sufficiently many cell divisions. This implies *β* > 1/3 for growing populations, if *α* was interpreted as random cell death. Computer simulations confirmed that for our purpose indeed *α* and *β* are interchangeable. Interestingly, this also suggests that in real cancer genomics data we should always find *β* > 1/3. Interestingly in 16 cancers analysed, in all cases we have *β* > 1/3, but we find two examples that are close to the predicted minimal possible value for growing populations (0.34 & 0.36).

### Non-constant cell death and tumour age inferences

Cell death is cell intrinsic in our model. This cell intrinsic death may have different underlying causes and the combined effect of these causes corresponds to the inferred *β* parameter. However, it is also possible that death varies with time, e.g. positive selection could select for lineages that avoid programmed cell death or escape the immune system more efficiently. To show the effects of such a change on the errors of our estimates one can for example think of the following time dependence of the cell-survival rate:

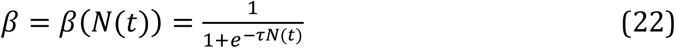

This Fermi-function is a particular but common choice to model such time dependences. Here, *N*(*t*) corresponds to the number of tumour cells at time *t* and *τ* is a free parameter that can be chosen to adjust the precise shape of the Fermi function. In this example, the function is such that at time *t* = 0 tumour cells start with a survival probability *β* = 1/2. Cell survival rates increase with time (positive selection) and will reach the maximum survival rate *β* = 1 at a certain tumour size *N* (SI Figure 24a). The parallel change of *β* for all cells simultaneously is unrealistic, but represents a worst-case scenario for our inference scheme.

The most critical implication of the parameter *β* is the estimation of tumour age. What error would we do, if such a process occurs undetected by our method? In this scenario average tumour growth is given by the differential equation:

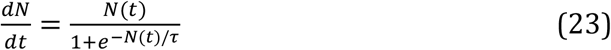

Assuming that a tumour is diagnosed at a certain size *N*_*D*_, then tumour age *T* is given (rearranging and integrating the above differential equation) by:

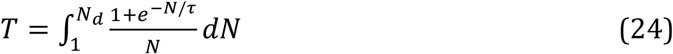

This integral cannot be expressed in simple analytical functions, but we can solve it numerically for any combination of *τ* and *N*_*D*_. Furthermore it is easy to see that the dominating term of the integral is of the order ~Log[*N*_*d*_]. To calculate the relative error, we would need to compare this time to the situation of unchanged *β*. This time is given by undisturbed exponential growth and is

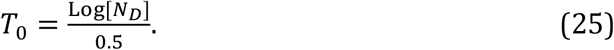

The relative error in the tumour age estimation then is 1 − *T*/*T*_0_. We can choose the scale parameter *τ* such that all tumour cells will have acquired the maximum possible per-cell survival rate *β* = 1 at a certain tumour size, e.g. for *τ* = 10^6^ cancer cells (at least 2 orders of magnitude below the current detection threshold). SI Figure 24b shows the relative errors of age inferences for tumours diagnosed at different sizes *N*_*D*_. The error increases for larger tumour size at diagnosis, as fitter cells have more time to cause deviations from the original prediction. However, even in the worst case scenario, the error remains < 20% and in more realistic situations is < 5%. Therefore, even in the worst-case scenario and a hypothetical tumour age of e.g. 5 years, an error of 20% adds an uncertainty of ±1 year. In most situations the deviation would be much smaller.

### Dependence of tumour age estimation on tumour size at diagnosis

Given a per-cell survival rate *β* we can estimate the number of generations necessary to generate a tumour of certain size. The time to diagnosis *T*_*D*_ only depends logarithmically on the number of tumour cells *N*_*D*_ at diagnosis

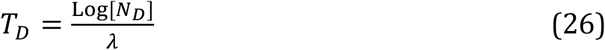

given some arbitrary proliferation rate *λ*. Again we can ask, what is the relative error if a tumour is diagnosed at a different size *N*_Δ_. As before, the relative error *η*_Δ_ is given by

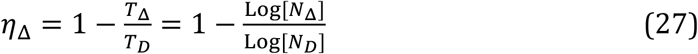

SI Figure 24 shows that even if the number of tumour cells differs by orders of magnitude, the relative error in tumour age estimations remains small. One order of magnitude discrepancy in the tumour size estimation corresponds to approx. 10% error of tumour age estimates.

### Individual based stochastic simulations of tumour growth

Individual based stochastic simulations of mutation accumulation in spatially growing tumours confirmed our premise *in silico*. We simulated tumours of ~1 million cells on a 2 dimensional grid with varying birth death and mutation rates using an implementation of the Gillespie algorithm (SI Figure 26). A cell division produces two surviving cells with probability *β* or one surviving cell with probability 1 − *β*. We also implemented an alternative version, where each daughter cell survives with a probability *α* after cell division. These simulations confirmed the equivalence of *α* and *β*. During each cell division, each daughter cell inherits the mutations of its parent and in addition accumulates novel mutations. The number of novel mutations is drawn from a *Poisson* distribution with mean *μ*. During simulations, the mutations for each cell as well as the division history of each cell are recorded.

We took bulk samples (between 100 and 10k cells per bulk) from each simulated tumour. These samples were either distributed at maximal distance (SI Figure 26a) or randomly distributed (SI Figure 26b). For most inferences, we used maximal distance sampling. Sequencing errors were simulated for each bulk by binomial sampling assuming sequencing depths of 100x or 200x, generating realistic mutation distributions comparable to available cancer genomic sequencing data. We then constructed all pairwise mutational distances of all ancestral cell lineages for each simulation separately. Figure 1 shows an example of the mutational distance distribution from 200 simulated tumour samples. It resembles the theoretically predicted mutational distance distribution almost perfectly.

To infer the mutation rate *μ* and the per-cell survival rate *β* solely from the distribution of mutational distances without any prior knowledge on the simulation, we adopted a probabilistic Bayesian Metropolis-Hastings (MCMC) algorithm (see also below for details). 9 bulk samples allow to infer the true per-cell mutation rate *μ* and per-cell survival rate *β* with high precision (*μ*: Spearman Rho = 0.98, *p* = 4×10^−23^; *β*: Spearman Rho = 0.93, *p* = 8×10^−16^, Relative error: *η*_*μ*_ = 0.056, *η*_*β*_ = 0.045), Figure 1e. The robustness of the inference scheme to more relaxed model assumptions and data quality and quantity are discussed below in more detail.

### Bayesian parameter inference

We now discuss the inverse problem, that is, can we reliably identify the single cell parameters (per-cell mutation rate, per-cell survival probability) given measured distances of multi-region sequenced cancer samples. These distances might be inferred from either forward in time simulated tumour growth, or data from multi-region sequenced cancer samples.

We use a Markov chain Monte Carlo method (MCMC), more precisely we implemented a standard Metropolis-Hastings-algorithm. The algorithm works as follows:

i. Create a new random set of model parameters ***w*** given the current set of parameters ***ν*** from a defined probability density *Q*, such that *Q*(*x*|*y*) = *Q*(*y*|*x*).
ii. Calculate the likelihood *L*(*P*(***w***)) of the model distribution *P*(***w***) given the data.
iii. Calculate the ratio of the new and old likelihood *ρ* = *L*(*P*(***w***)/*L*(*P*(***ν***)). Accept the new parameter set with probability *ρ* otherwise reject.
iv. Repeat

In our case the model distribution is given by equation (13). To calculate the likelihood of equation (13) given the data, we have to choose a cut off for the two infinite sums. However, real data always has a maximum mutational distance. Higher terms of the infinite sums contribute to higher mutational distances. The distribution of interest does not change for a sufficiently high cut off and each observed data set only requires finite many terms. We used uninformed uniform prior distributions for the per-cell mutation rate *μ* and the per-cell survival rate *β* in all cases. Point estimates were extracted as sample medians from the MCMC inferences for the mutation rate and cell survival rate separately. The ranges of the uniform priors can be adjusted to optimise acceptance rates and computational time. In our implementation, a new set of parameters is always relative to the previously accepted parameter set ***w***_New_ = ***w***_*Old*_ + Φ(***w***), where Φ is the prior parameter distribution. A typical range used in our inference scheme is Φ_uniform_(*β*) = [−0.06, +0.06] and Φ_uniform_(*μ*) = [−5, +5] for whole genome sequencing, but could also vary for data sets with higher or lower mutational burden. We tested the robustness of the MCMC framework and in addition used Gamma distributions as prior (SI Figure 27). The MCMC inference converges to the same parameter sets, independent of the prior distribution of choice, or the initial starting condition of the Markov Chain. If both parameters are inferred simultaneously, they converge to the correct initially set of parameter values. If one of the parameters is fixed to the correct value, the second parameter does converge to the correct value (SI Figure 27).

Examples of MCMC chains and resulting parameter distributions are shown in SI Figures 1–3, SI Figures 10–18 & Si Figures 27, 28. Our method is able to recover the parameter sets of simulated tumour growth reliably with high precision, see Figure 1 in the main text, e. g., 9 bulk samples allow to infer the true mutation rate *μ* and per-cell survival rate *β* with high precision (*μ*: Spearman Rho = 0.98, *p* = 4×10^−23^; *β*: Spearman Rho = 0.93, *p* = 8×10^−16^, Relative error: *η*_*μ*_ = 0.056, *η*_*β*_ = 0.045). Inferences remain possible for up to 6 independent bulk samples (SI Figure 33) and reduced sequencing depth of 25x (SI Figure 32).

### Robustness of evolutionary parameter inferences on model assumptions and data quality and quantity

Our theoretical derivation of the mutational distance distribution and our computational framework of per-cell mutation and per-cell survival inference are based on a set of assumptions. In the following we discuss these assumptions in more detail and quantify the robustness of our parameter inferences to violations of these assumptions.

### Non-constant per-cell mutation rate

We assume that with each cell division cancer cells acquire *X* novel mutations, where *X* is a Poisson distributed random variable with a constant mean mutation rate *μ*. However, in principal, the mean mutation rate *μ* could itself vary. These changes might occur in bursts (e.g. sudden APOBEC activity) or change more steadily over time. We discuss two additional scenarios. The first scenario allows the rate *μ* to be a random variable. We choose an exponential distribution *P*(*μ*_*Exp*_) = (1/*λ*)*e*^*−μ/λ*^. In this scenario one allows for considerable noise in single cell divisions, for example cell divisions with lower mutation rates and a few cell divisions with much higher mutation rates (modelling e.g. random bursts due to APOBEC activity or other events). In the second scenario we can consider a situation where the mutation rate grows with time. e.g.: *μ*(*t*) = *μ*_0_ + 0.5Log[2]×*t*. For our simulations, this implies an approximately 10-20 times increased mutation rate at the time of sampling compared to the beginning.

Parameter inferences for these scenarios are shown in SI Figure 28. An exponentially distributed rate parameter *μ* (the number of mutations remains Poisson but with the non-constant rate parameter *μ*) adds noise to the distribution of the mutational distances compared to the constant rate scenario (e.g. SI Figure 28a-f), but the parameter inference remains robust (e.g. SI Figure 28g-l), although leading to slightly lower estimates of survival rates. The second scenario of a linear increasing mutation rate causes significant error in the estimation of the mutation rate parameters (SI Figure 28m-r), mostly because of the distortion of the mutational distance distribution, against which the theoretical model does not fit (SI Figure 28s). However, the inferred effective mutation rate corresponds to the time averaged mean mutation rate on the course of the stochastic simulation. In SI Figure 28t,u we show the relative error in the estimation of the parameters under the different mutation rate models (20 instances of simulated tumours per scenario).

### Non-exponential tumour growth

The analytical derivation of the mutational distance distribution is based on exponentially growing tumour. Although space is not modelled explicitly in our theoretical derivation, it is an essential ingredient as only spatial sampling allows constructing sufficiently many different cell lineages within single tumours. The mode of growth, e.g. exponential vs. peripheral could alter the properties of which and how many different lineages are sampled in space. We therefore introduced an additional parameter *a* into our computational stochastic tumour simulations that models the ability of a cell to proliferate in the presence/absence of empty space in its direct neighbourhood. If *a* = 1, a cell can always proliferate regardless of empty space in its direct neighbourhood by pushing cells and creating an empty spot. This results in exponential growth (SI Figure 29a & d), however the spatial proximity of more recent offspring is maintained. If 0 ≤ *a* < 1 cells only proliferate with a probability according to the value *a* in the absence of empty space, thus favouring cells at the less dense peripheral boundary of the tumour. This leads to polynomial growth (SI Figure29c & f). We tested the ability of our method to robustly recover the mutation rate for different values of *a*, such as *a* = 1 (SI Figure 29a), *a* = 0.5 (SI Figure 27b) and *a* = 0.25 (SI Figure 29c). Although the variance of the parameter inference slightly increases, the relative error *η* remained below 10% in the vast majority of cases (SI Figure 29). Hence, deviations from exponential growth increase uncertainties, but the inference remains robust.

### Partial selective sweeps

Positive selection and partial selective sweeps could impact our inference of evolutionary parameters. In order to test the robustness of our method to the effects of selection, we ran simulations with varying strength of positively selected clones. Here positive selection confers a proliferation advantage to cells. SI Figure 26 shows examples of 10 simulated tumours as well as the spatial sampling schemes, such as maximum distance spatial sampling (SI Figure 26a) vs. random spatial sampling (SI Figure 26b). We selected simulations with a partial selective sweep and excluded scenarios where the selected subclone reached fixation before sampling (in this scenario the tumour goes back to be uni-clonal and thus within-clone neutral again). SI Figure 30 provides a summary of the inferences for neutral tumours (selection coefficient s=0; SI Figure 30a), compared to tumours with varying selection strength (s=0.1, s=0.25 and s=0.5; SI Figure 30a & b). Note the high coefficients of selection that infer fitness advantages of up to 50% (s=0.5) to a selected sub-clone in our simulations. The presence of positively selected sub-clones adds uncertainty to the mutational distance distributions. However, even for very strong positive selection of *s* = 0.5 (corresponding to a 50% fitness advantage) mutation rate inferences remain robust (relative error: *η*_*μ*_ = 0.1) and also per-cell survival rate estimates remain stable (relative error: *η*_*β*_ = 0.1 for *s* = 0.5, SI Figure 30).

### Spatial sampling strategies and sampling biases

The construction of mutational distances relies on detecting differences between ancestral cells that are inferred from mutations of bulk or single cell samples. One would expect that genetic and spatial distance of samples is on average positively correlated (although the relation is non-linear and probably more involved)^34^. Thus the relative location of samples potentially influences inferences. We compared parameter inferences from tumours with a maximal distance sampling strategy (which is the most common current sampling strategy in clinical practice) and a random sampling strategy, where positions of sampled bulks are assigned randomly (SI Figure 26). In both cases, inferences are robust against different spatial sampling schemes for both the mutation rate (SI Figure 31a) and the survival rate (SI Figure 31b). Maximal distance sampling appears to perform slightly better compared to random sampling strategies. Inferences throughout this manuscript relied on a maximal distance sampling strategy, if not stated otherwise.

### Sequencing depth

Our stochastic simulations allow us to reproduce the effects of sequencing coverage on the final data and consequently our evolutionary parameter inferences. Briefly, we generate dispersed coverage values for input mutations. We do that by sampling a coverage from a Poisson distribution: Poisson (*λ* = Z) with mean *λ* equal to a desired sequencing depth *Z*. Once we have sampled a depth value k for a mutation, we sample its frequency (number of reads with the variant allele frequency) with a Binomial trail. We use *f* ~ Binomial(*n, k*), where *n* is the proportion of cells carrying this mutation given all cells sampled in the simulated biopsy.

In SI Figure 32 we show that parameter inferences based on the mutational distance distribution remain robust for low sequencing coverage. Sequencing depth in the available data varied. In general Exome sequencing had very high coverage: Lung Cancer TracerX median coverage ^28^: 426x, Renal cancer analysis coverage ^29^: >100x. Whole genome sequencing had lower coverage: Median coverage of colorectal cancers from Cross et al. ^21^: 55x. Whole genomes of single colon stem cells in Roerink et al. ^27^ were sequenced from single cell derived Organoids sequenced at 30x coverage.

### Number of independent samples per tumour

SI Figure 33 shows parameter inferences for multiple independent spatial tumour simulations with 6 to 9 bulk samples. Fewer samples, as expected, increase the noise of the mutational distance distribution. Parameter inferences from up to 6 samples remain robust. In the analysis of real tumours in this manuscript, the distribution of independent samples per tumour is (*number of cases* × *number of independent samples*): 2×6; 4×*7*; 5×8; 4×9; 1×13. This corresponds to 129 tumour samples in total with a median of 8 samples per tumour. In addition, we also include the analysis of 89 whole genome sequenced healthy haematopoietic stem cells from a single healthy donor (Figure 2 in the main text).

### Genomic analysis of cancer samples

Details of the bioinformatics-analysis of the multi-region sequenced tumour samples can be found for the colon carcinomas and the adenoma in^21^, the additional three single cell whole genome sequenced colon cancers^27^ the renal cell carcinomas in^29^ and the lung squamous and adenocarcinoma in^28^. Details on the methodology and sequencing of single stem cells in healthy haematopoiesis can be found in^20^.

### Mutational signature inference

For each sample we found the set of signatures (among those signatures reported in CRC) that best explained the totality of mutations in the sample. Specifically we did a non-negative regression of the sample’s mutations against all the CRC signatures^24^ and found those signatures with non-zero coefficients. We took these as the candidate signatures for each sample.

For each mutation in each sample, we determined the likelihood of the mutation under each of the candidate signatures. We assigned a mutation to a candidate signature where the likelihood under that signature was at least twice that under any other. If there was no such signature, we assigned the mutation to ‘Signature.Other’. The method was originally developed in^35^ and is based on the R-package “SomaticSignatures” ^36^. We did not adjust for differences in nucleotide composition when calculating differences between coding and non-coding regions as we wanted to infer the overall point mutation rate in these regions, see for example SI Figure 5. Nucleotide dependent mutation rate estimates are shown in SI Figure 6. Nucleotide composition was adjusted for to calculate the mutation rates of mutational signatures using standard tools^36^.

### Per-cell survival rate and stem cell properties

The data from Hans Clevers and colleagues^27^ measures mutational burden in single colon stem cells by expanding isolated donor derived single cells into organoids. Thus in this case the inferences most certainly correspond to stem cell population dynamics. Furthermore, our analysis relies on mutational distances between ancestral cells and thus surviving lineages within the tumour population throughout its evolutionary history. These lineages of ancestral cells probably also represent stem cell lineages. If there is a dichotomy of stem and non-stem cells in these tumours, our method corresponds to the survival rate of stem cell lineages. This is further supported by the fact that our model describes the mutation accumulation in healthy haematopoietic stem cell lineages well (Figure 2 in the main text). Ultimately, it is the expansion of the stem cell population that determines tumour growth. In particular for solid tumours there is accumulating evidence that the fraction of stem cells is high compared to most blood cancers, explaining the common failure of targeted therapy in solid tumours^37,38^.

## Supplemental Figures

**SI Figure 1:**
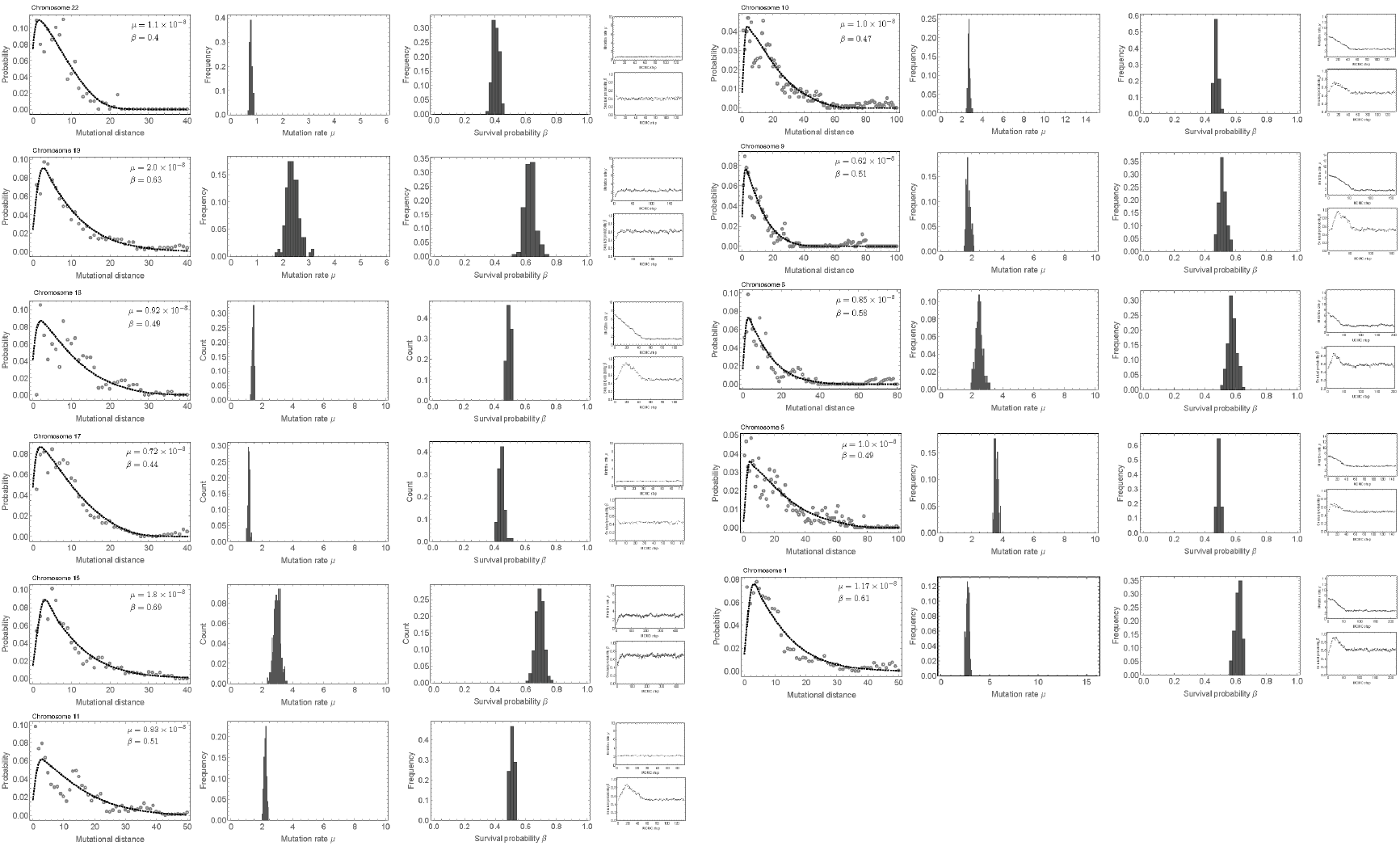
Patient 02 mutational distance distribution and MCMC parameter inference per chromosome. Chromosomes with sub-clonal copy number alterations were discarded from the analysis. Original data taken from^21^.

**SI Figure 2:**
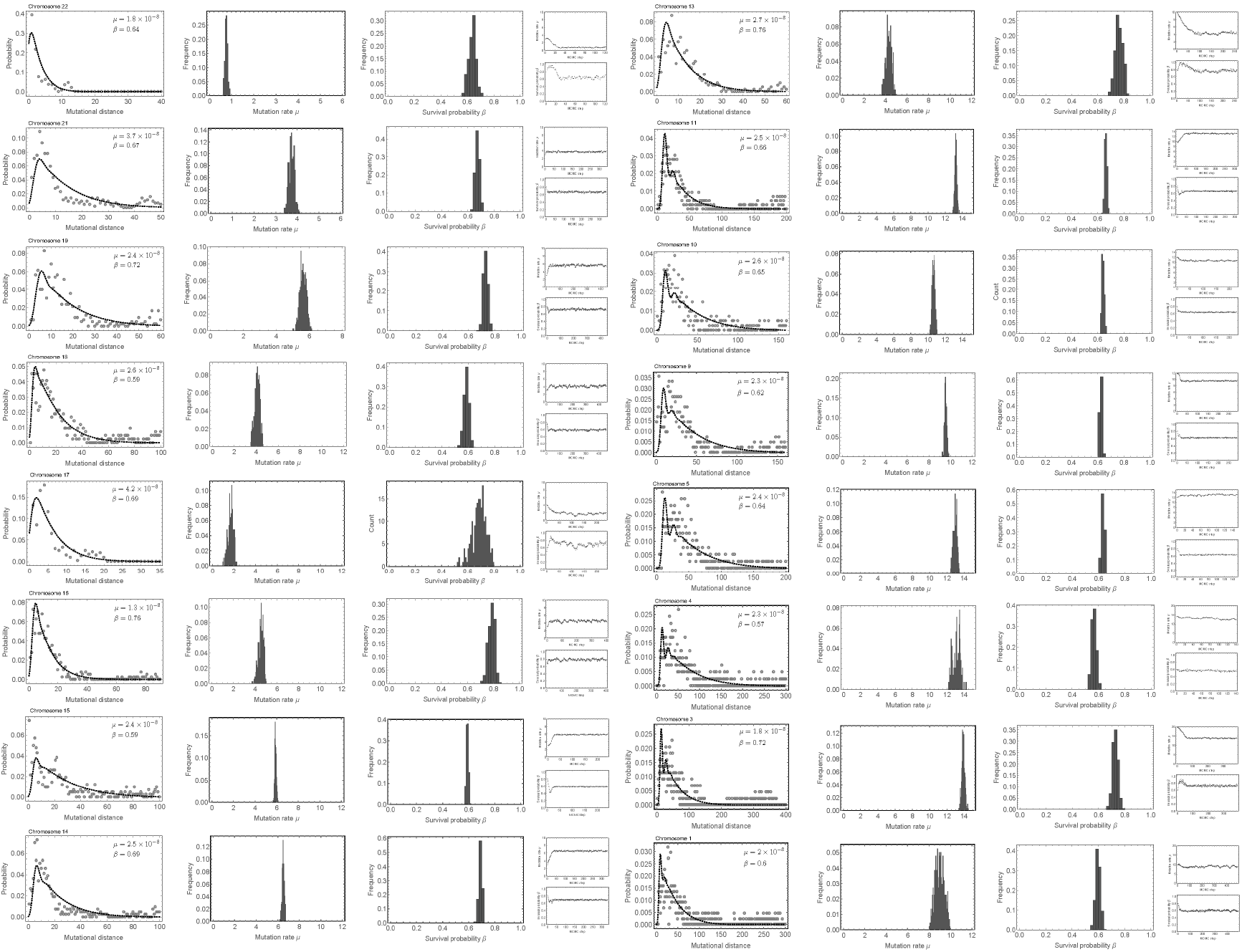
Patient 03 mutational distance distribution and MCMC parameter inference per chromosome. Chromosomes with sub-clonal copy number alterations were discarded from the analysis. Original data taken from^21^.

**SI Figure 3:**
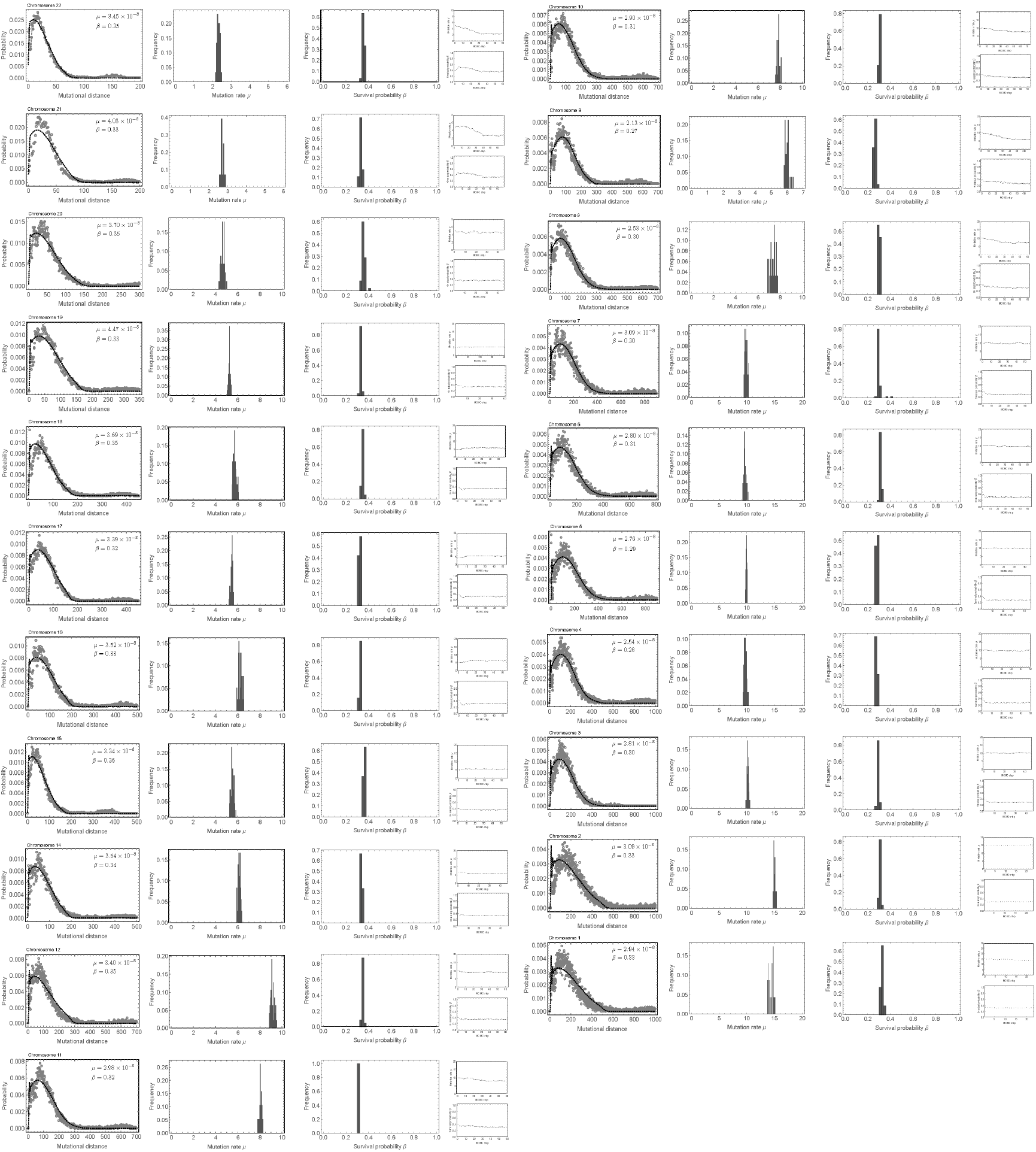
Patient 04 mutational distance distribution and MCMC parameter inference per chromosome. Chromosomes with sub-clonal copy number alterations were discarded from the analysis. Original data taken from^21^.

**SI Figure 4:**
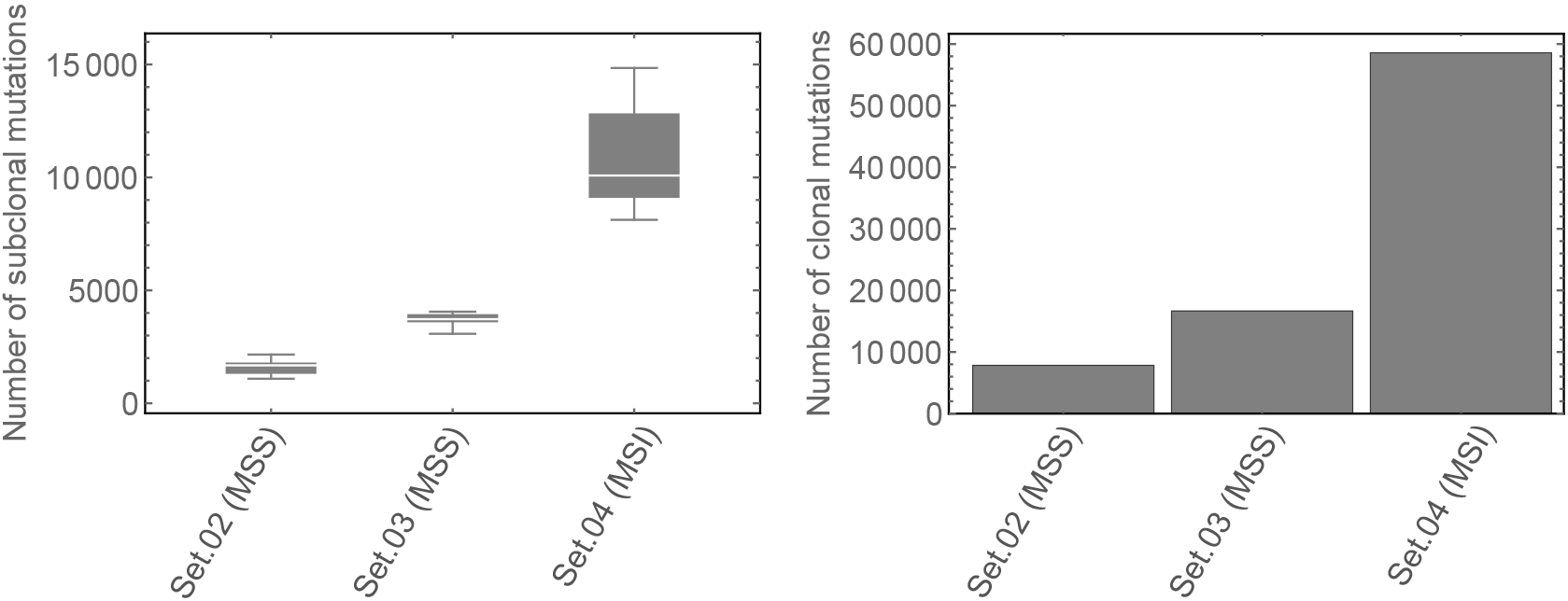
Mutational load of sub-clonal and clonal mutations of Patients 02-04. Mutations were classified as clonal if they were present in all bulk samples of the tumour^39^. Mutations present within a subset of samples of a tumour were classified as sub-clonal. MSI tumour 04 shows both higher clonal and sub-clonal mutational burden compared to MSS tumours 02 and 03.

**SI Figure 5:**
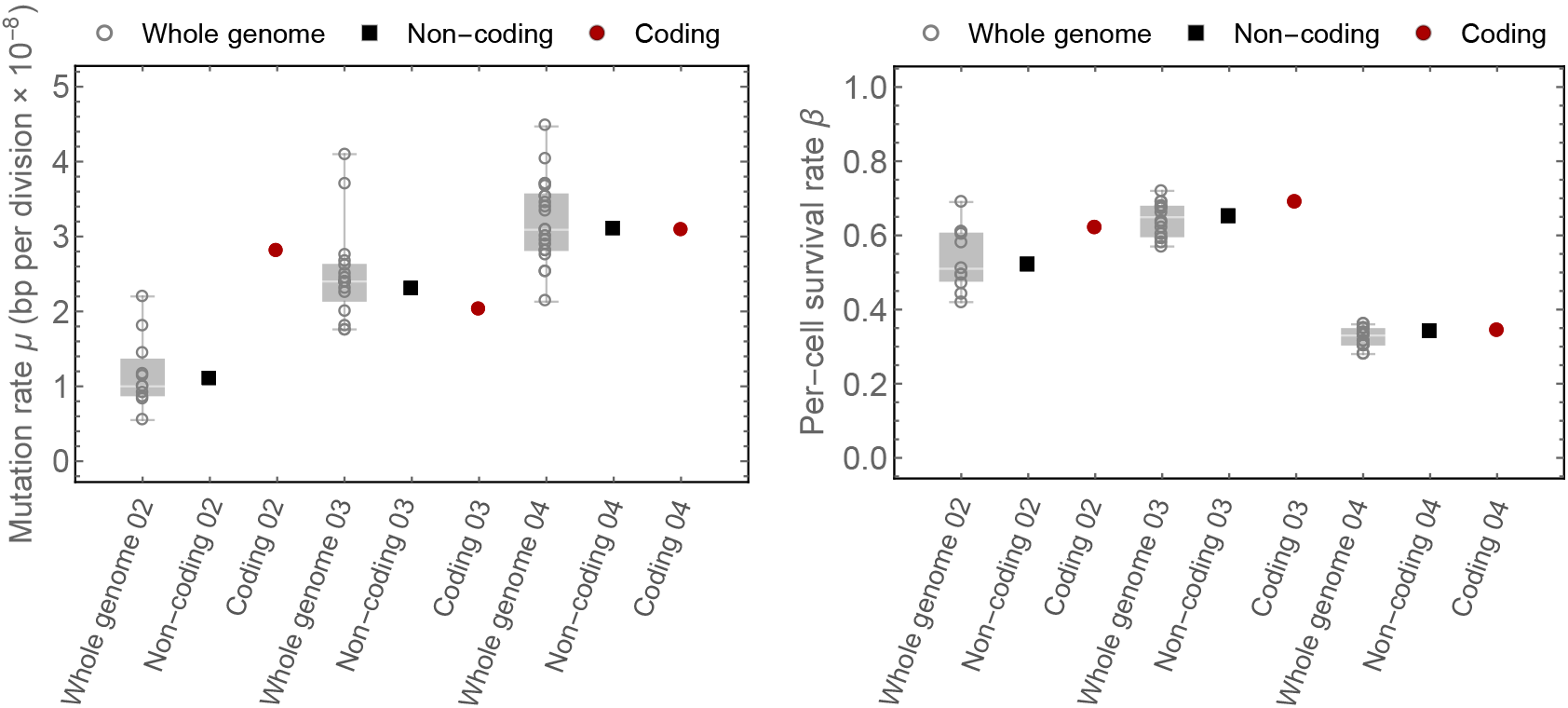
Inference of per-cell mutation and per-cell survival rate for whole genome (per chromosome, open grey circles), non-coding (black squares) and coding mutations (red circles) in Patients 02-04. The coding mutation rate in patient 02 is slightly increased compared to whole genome inferences 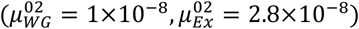, they are slightly lower in patient 03 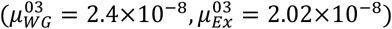 and the same in patient 04 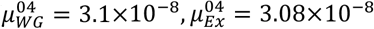. Non-coding mutation rates agree with median whole genome mutation rates. Recently it was suggested that mismatch repair efficacy differs in coding and non-coding regions of the genome^22^. Consistent with this hypothesis the MSI+ tumour in Patient 04 shows the exact same mutation rate per cell division in whole genome, non-coding and coding genome regions. The per-cell survival rate inferences are across whole genome, non-coding and coding genome regions.

**SI Figure 6:**
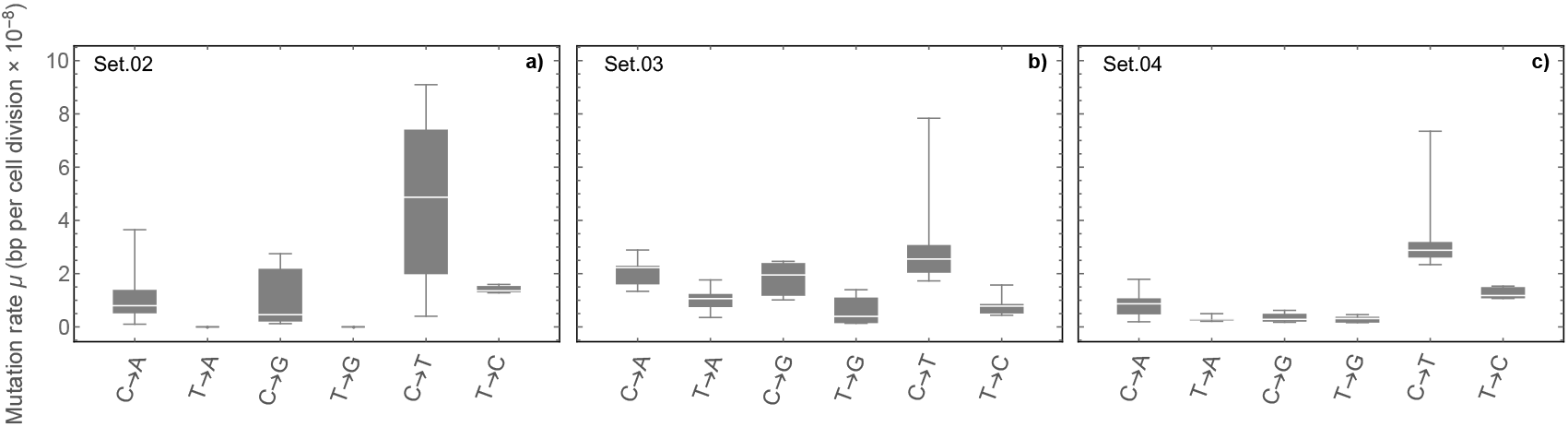
Mutation rates for mutational subtypes. The mutation rate for each mutational subtype was inferred based on our MCMC algorithm for individual chromosomes (see Figure 2 in the main text) for all 3 patients separately and normalised for the C & T content at each chromosome. In Patient 02 and 04 transitions show higher mutation rates than transversions. Interestingly, in Patient 02 the mutation rates for the transversions T → A and T → G are below a detection threshold, whereas in Patient 04 they are detectable. The overall pattern of somatic mutation accumulation in theses two patients agrees with patterns of genome divergence between human and chimpanzees26. Patient 03 shows a distinct pattern of mutation accumulation. Here transitions and transversions appear equally likely, with C → X mutations slightly more likely compared to T → X mutations.

**SI Figure 7:**
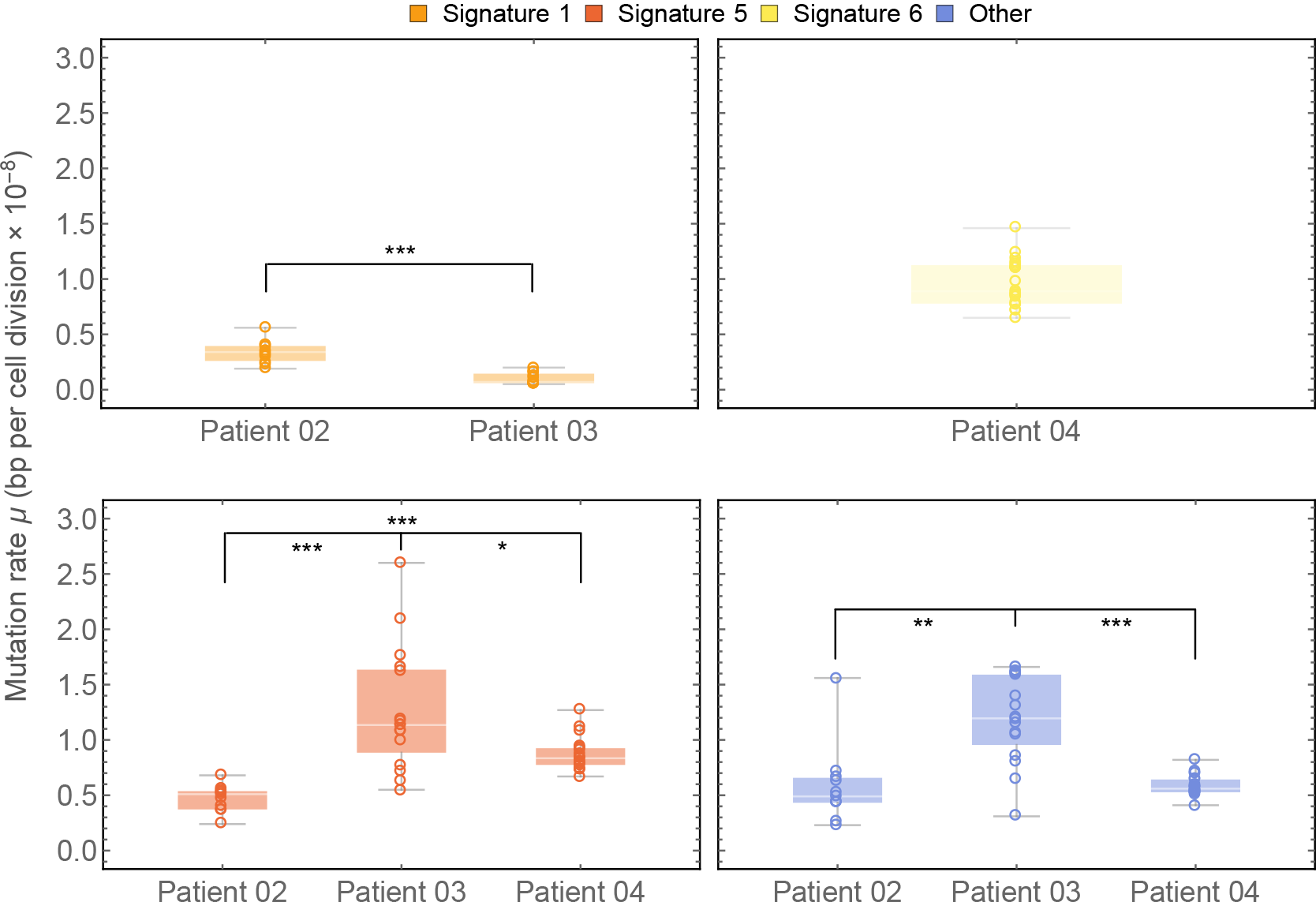
Distribution of mutational signature mutation rate per chromosome for Patients 02-04. Mutation rates per cell division of mutational signatures differ significantly between patients. (*: p<0.05, ** :p<0.01, ***: p<0.001, Mann-Whitney-U-test).

**SI Figure 8:**
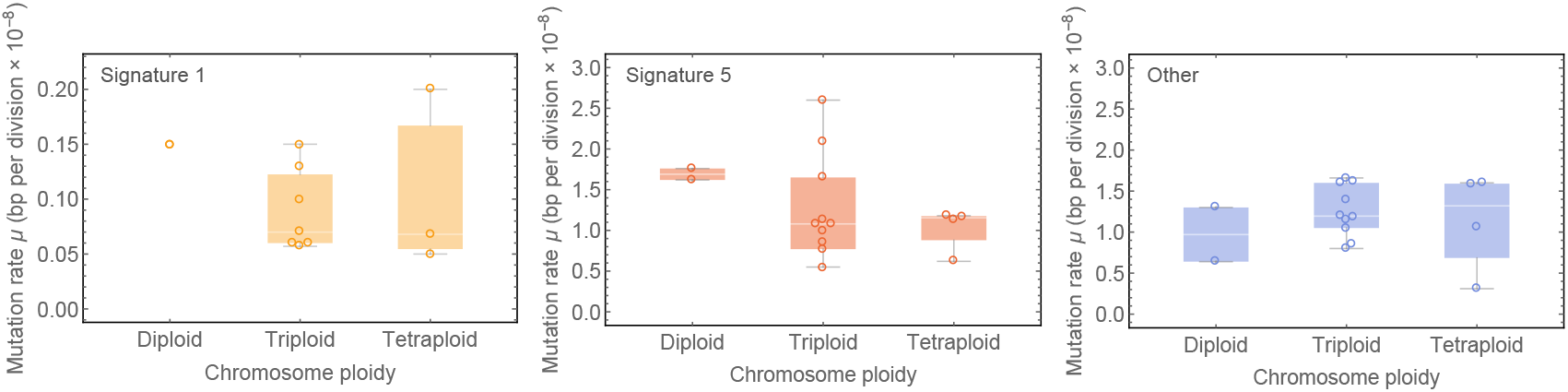
Distribution of mutational signature mutation rate for different chromosomes of Patient 03 sorted by ploidy. After normalizing for ploidy, mutation rate distributions do not differ significantly (p>0.3, Mann-Whitney-U-test).

**SI Figure 9:**
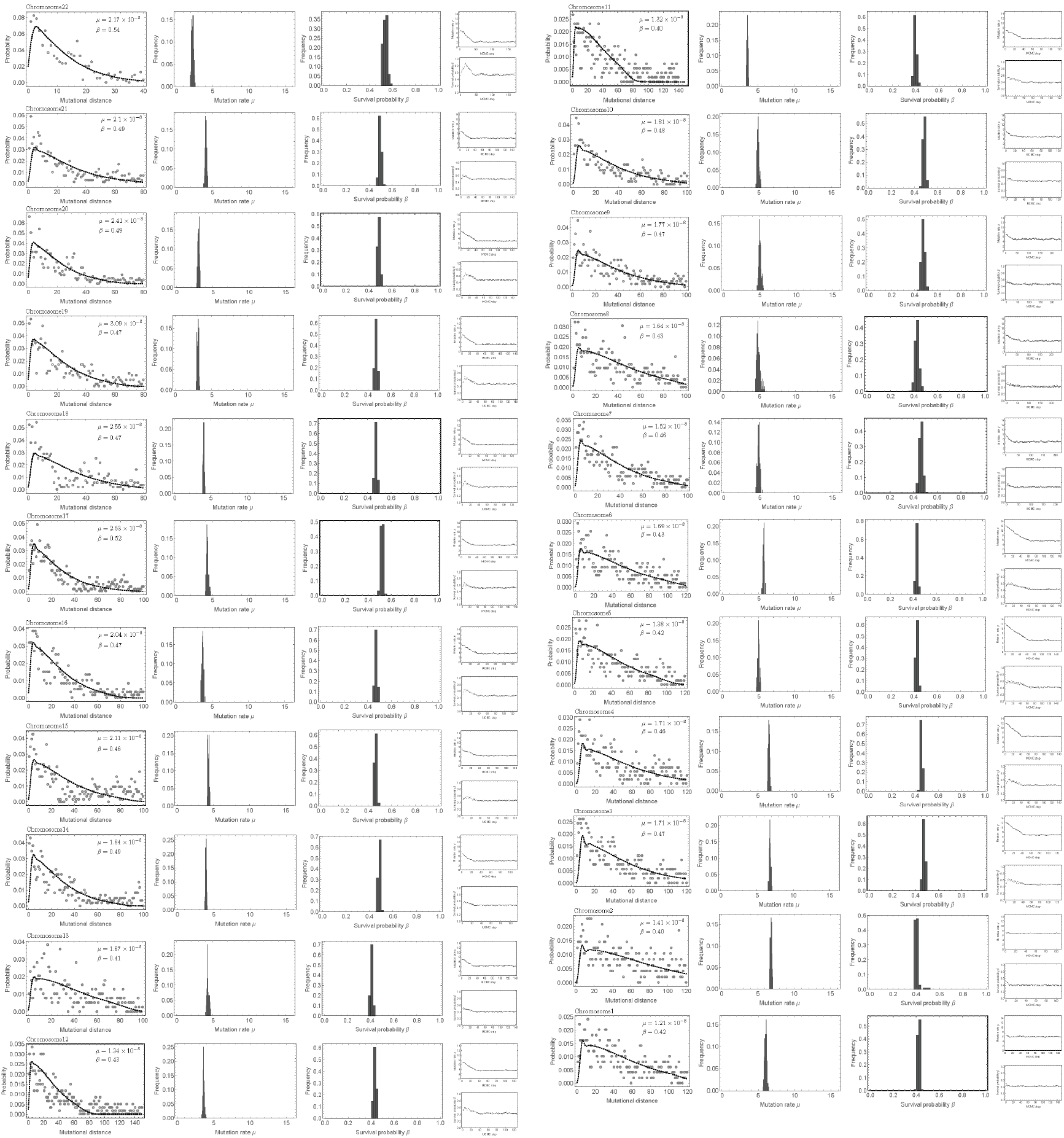
Mutational distance distribution and MCMC inference for individual chromosomes inferred from 7 whole genome sequenced samples of a MSI+ colon cancer patient. Data was taken from^27^

**SI Figure 10:**
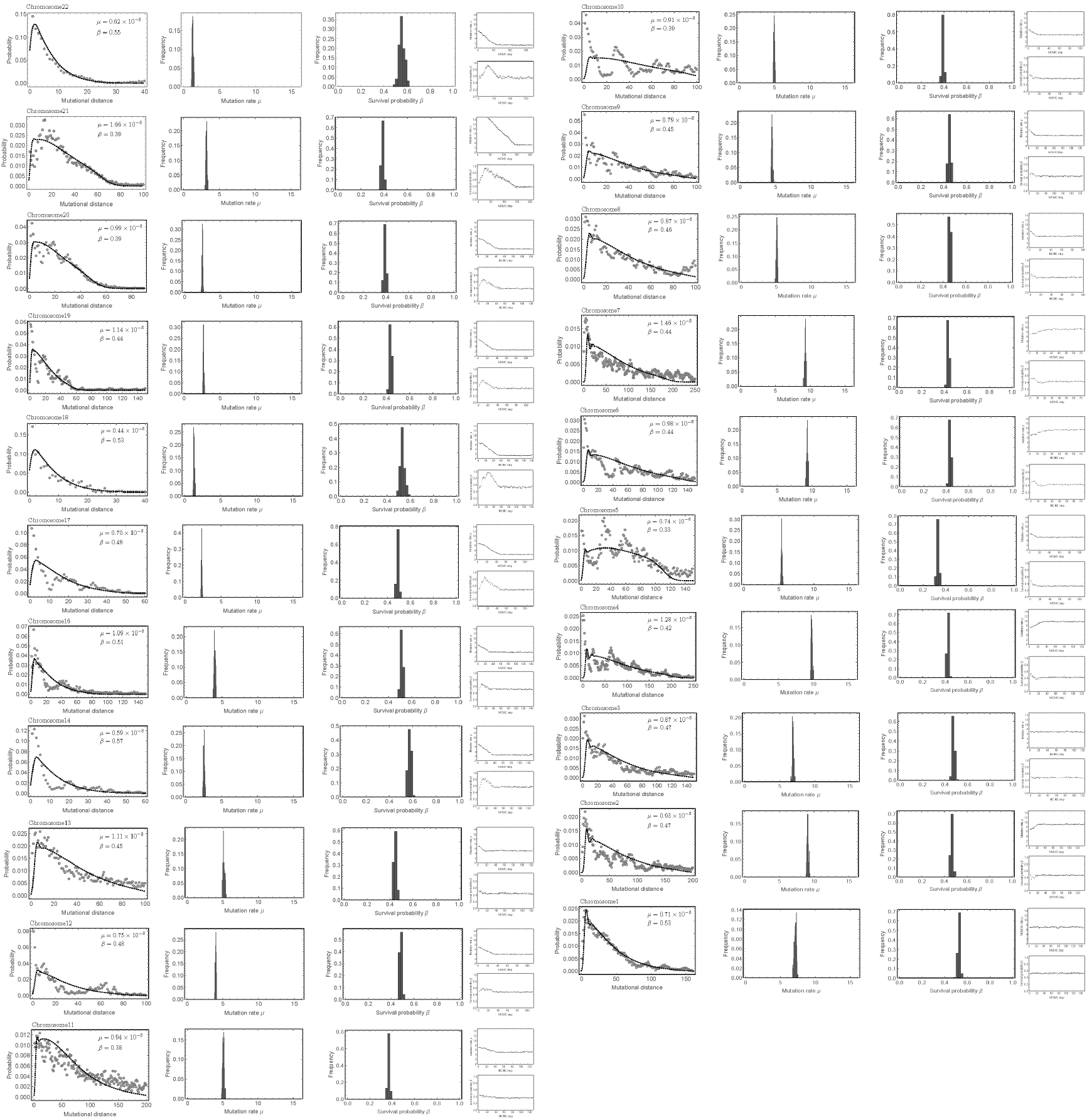
Mutational distance distribution and MCMC inference for individual chromosomes inferred from 9 whole genome sequenced samples of a MSS colon cancer patient. Data was taken from^27^.

**SI Figure 11:**
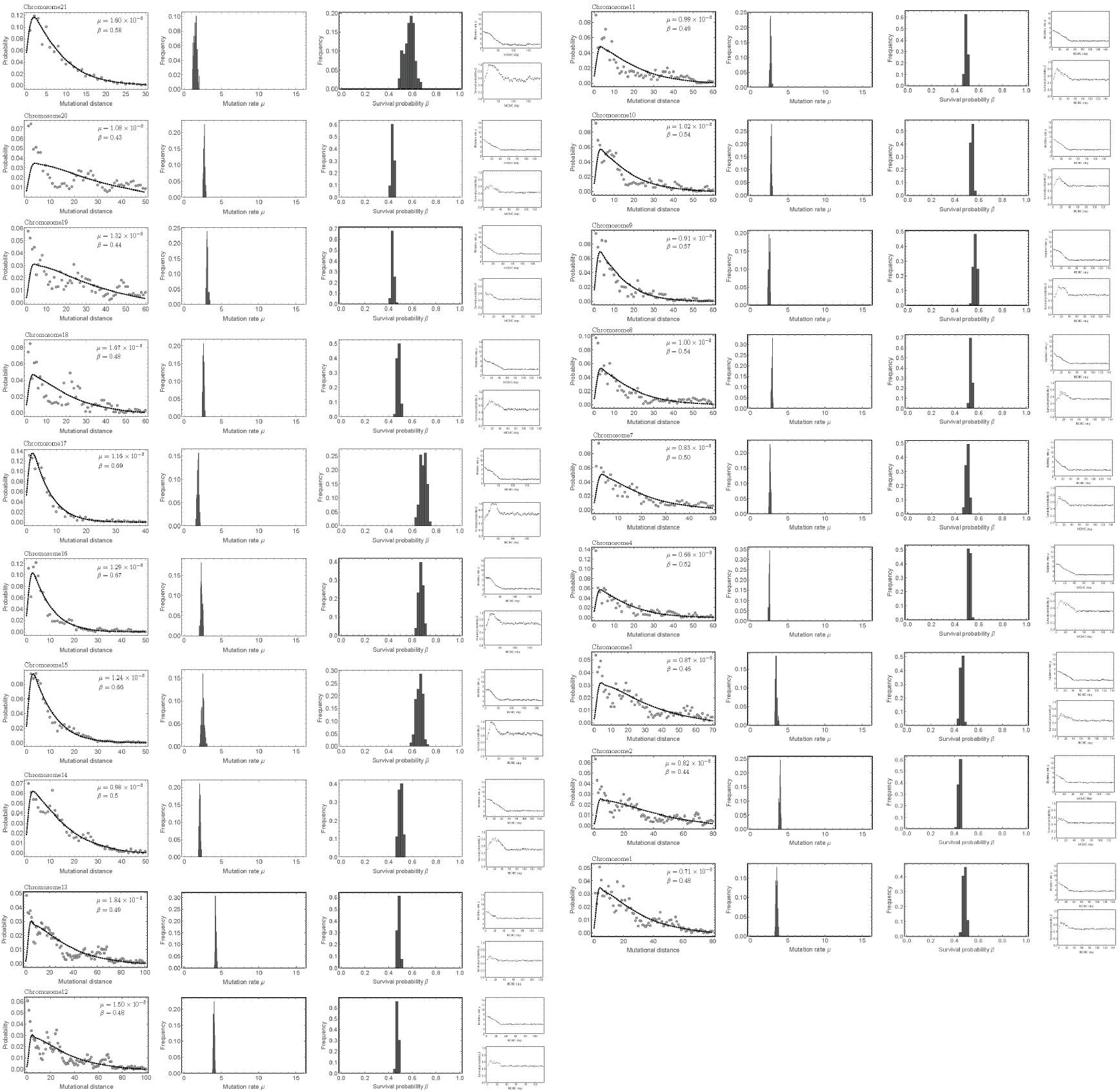
Mutational distance distribution and MCMC inference for individual chromosomes inferred from 9 whole genome sequenced samples of a MSS colon cancer patient. Data was taken from^27^.

**SI Figure 12:**
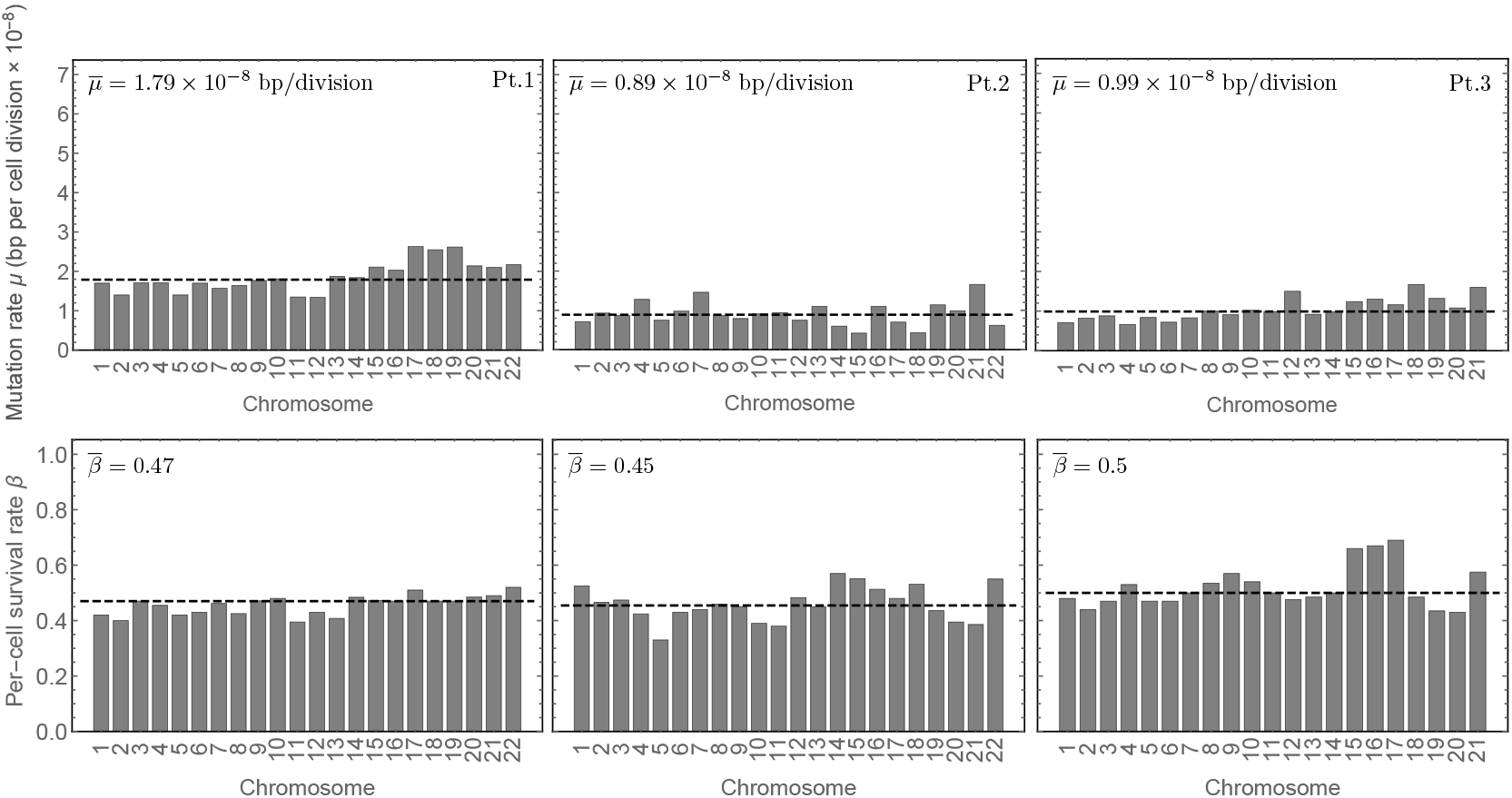
Inference of the mutation rate *μ* per cell division and the per-cell survival rate *β* per chromosome for the three patients shown in SI Figures 9–11. Insets show median mutation and per-cell survival rates. Patient 1 is MSI+ and presents with a higher mutation rate per cell division compared to patients 2 and 3. Data was taken from^27^.

**SI Figure 13:**
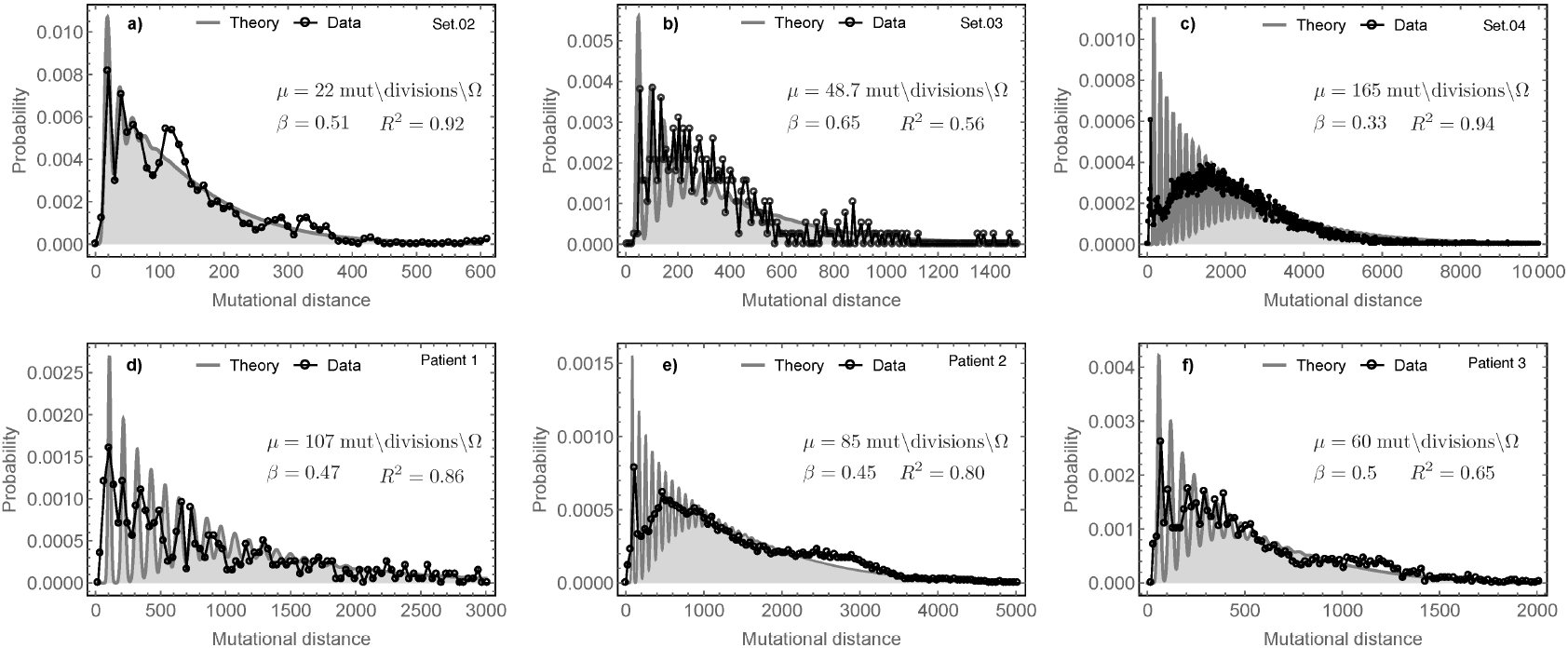
Mutational distance distributions constructed from the whole genome of a)-c) tumours 02-04 from^21^ and d)-f) Patients 1-3 from^27^. Grey lines show the theoretical mutational distance distribution based on best inferences and black dots correspond the mutational distances in the tumours. Note the different scales for the mutational distances and the order of magnitude difference between overall probabilities of distances across patients. These mutational distance distributions fall into different classes based on the per-cell mutation rate and per-cell survival rate as illustrated in SI Figure21.

**SI Figure 14:**
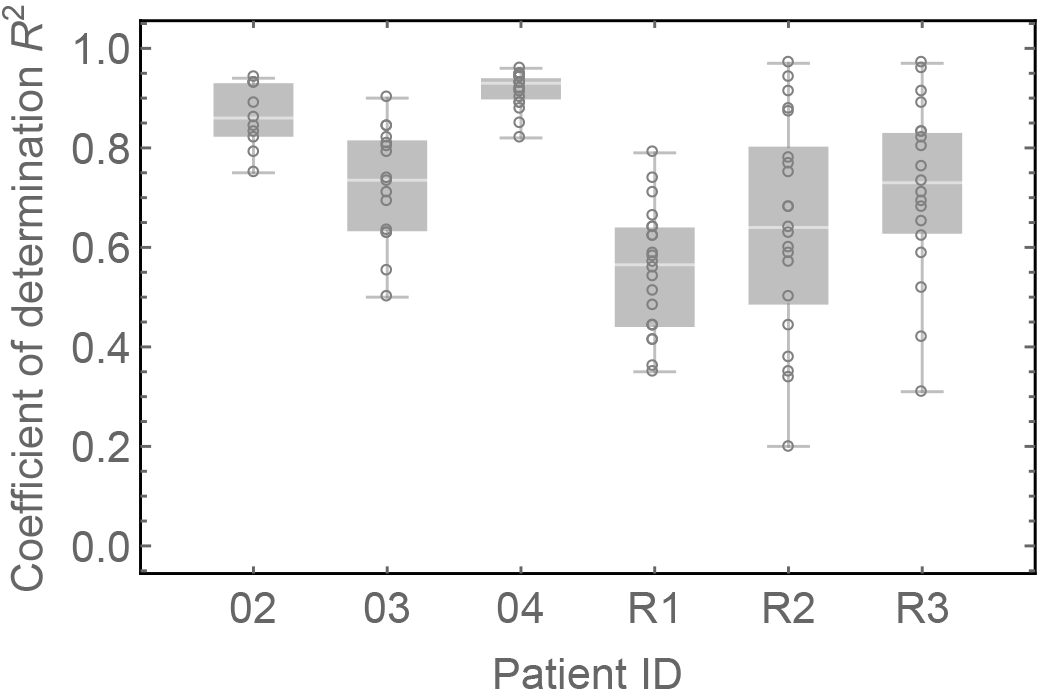
Coefficient of determination between per-chromosome estimates and the best estimates of the mutational distance distributions. Shown are the coefficients of determination per chromosome for tumours 02-04 and Patients 1-3 as shown in SI Figures 1–3 & 9–11.

**SI Figure 15:**
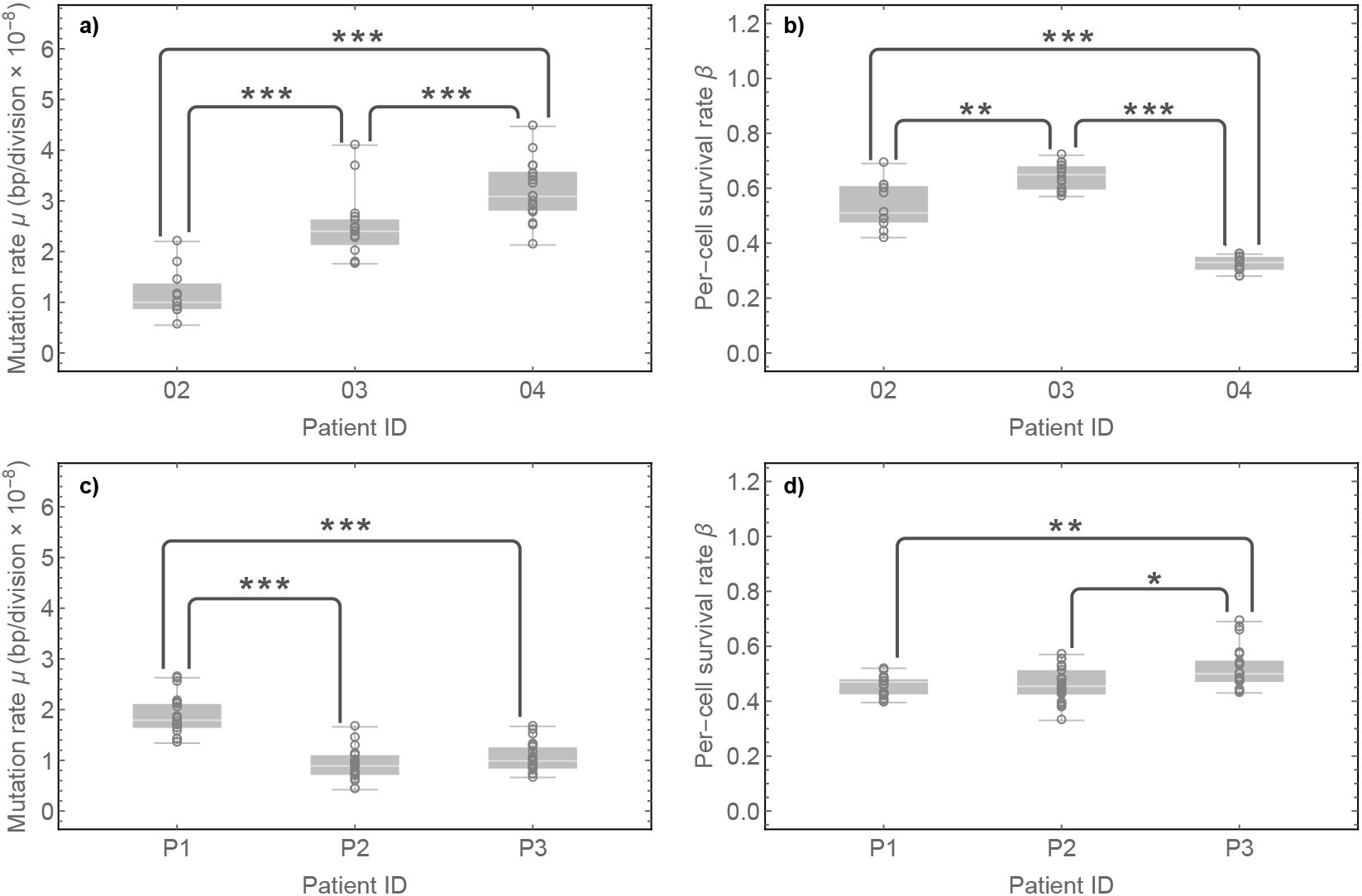
Between patient differences of evolutionary parameters. Shown are the inferences of the mutation and per-cell survival rates per chromosome (dots) for 6 whole genome sequenced colorectal cancers. Panels **(a)** and **(b)** show the cohort of Cross et al. 21 and panels **(c)** and **(d)** the original patients by Roerink et al.^27^ A Mann-Whitney-U-test was used to test between patient differences, symbols correspond to the short notation: * = *p* < 0.01, ** = *p* < 0.001 and *** = *p* < 0.0001.

**SI Figure 16:**
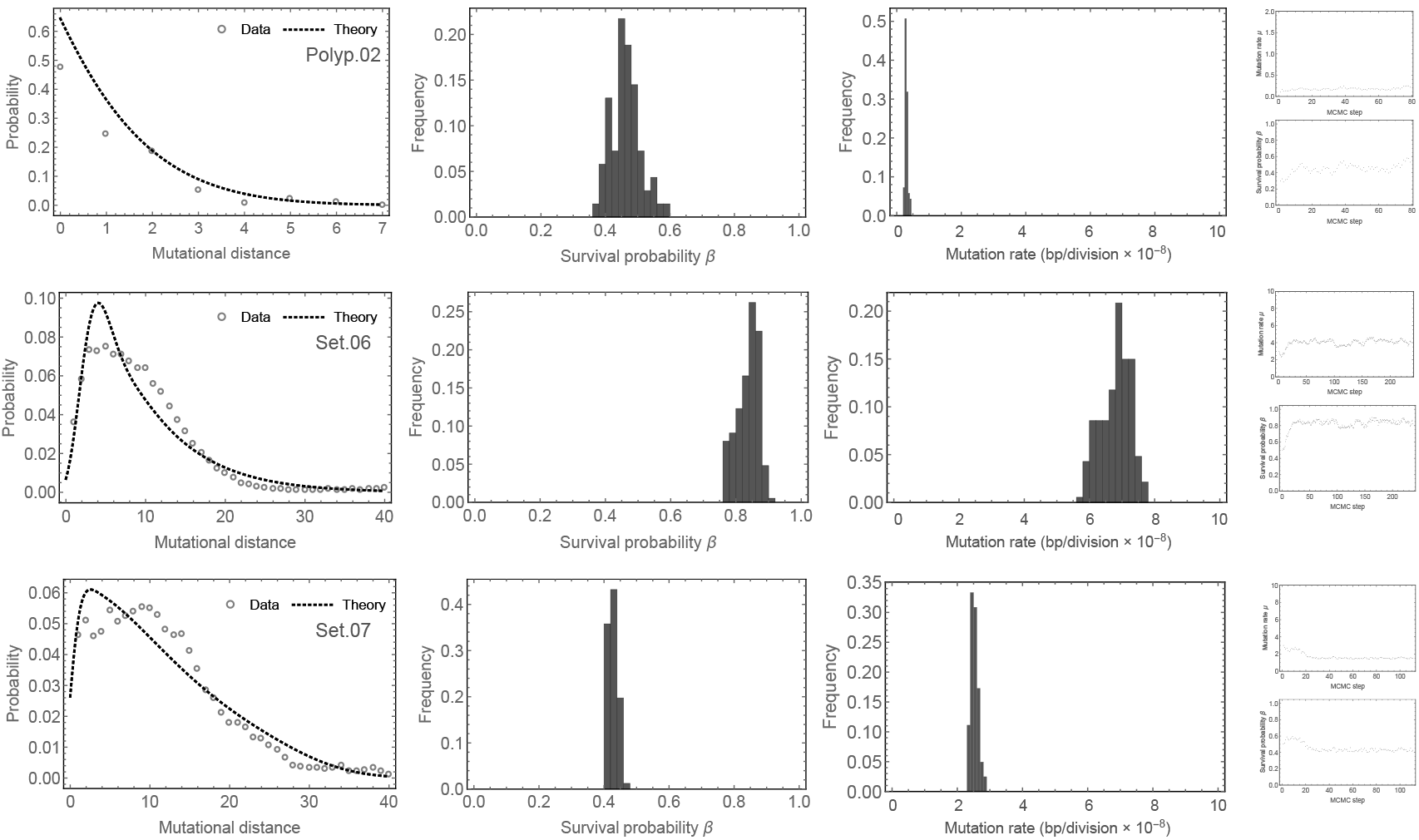
Mutational distance distribution and MCMC inference from a single exome sequenced adenoma (top) and two exome sequenced carcinomas (bottom). Note, the adenoma shows smaller mutational differences between ancestral cells compared to both carcinomas and presents with a near normal mutation rate, see main text also. However the per-cell survival probability is higher compared to normal tissue and the adenoma is expected to clonally expand.

**SI Figure 17:**
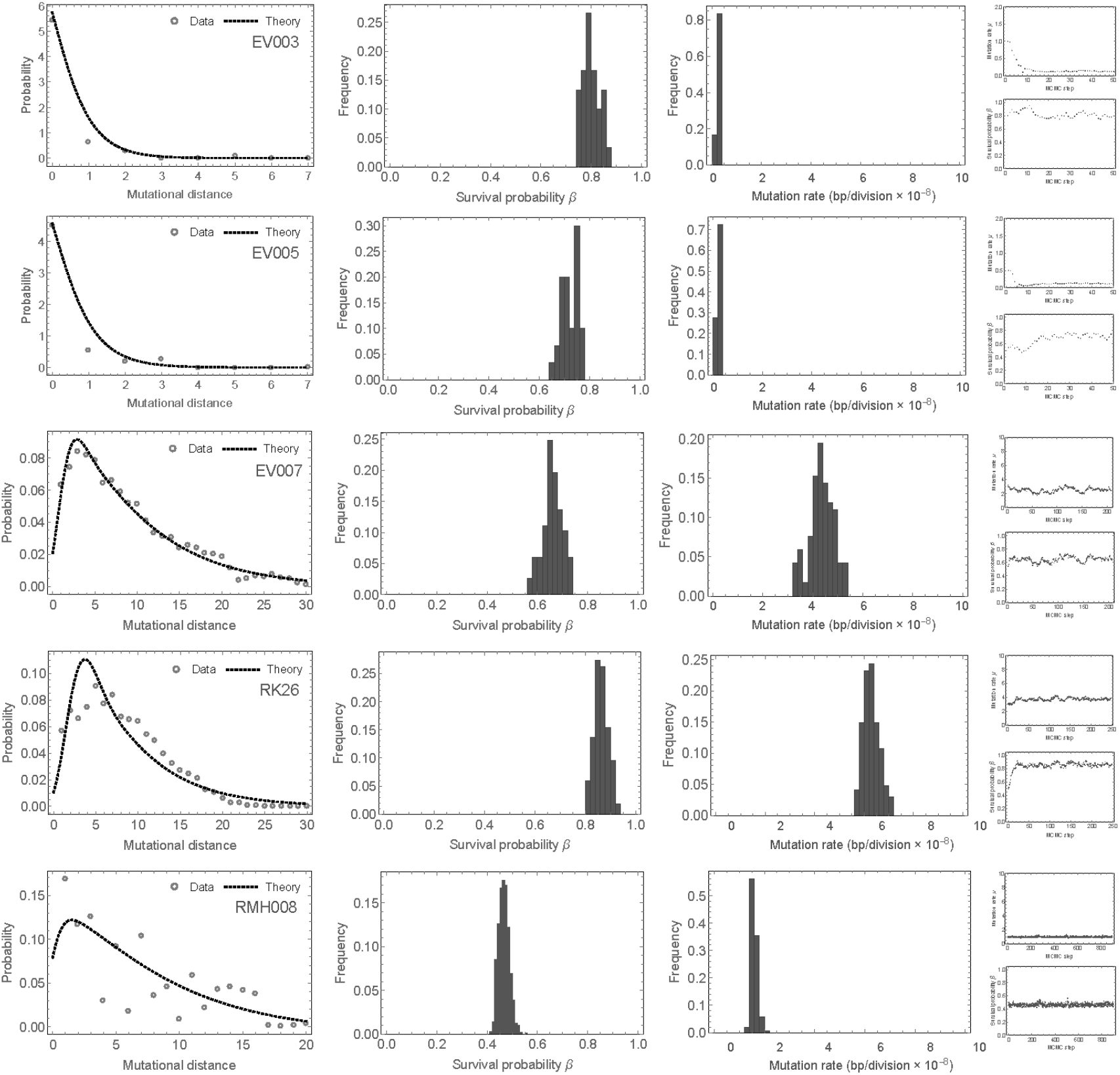
Mutational distance distribution and MCMC inference from 5 exome sequenced renal cell carcinomas^29^. Surprisingly, two renal cell carcinomas appear to have near normal mutation rates (EV003 and EV005), similar to the colon adenoma. However, all 5 cases present with high per cell survival probabilities.

**SI Figure 18:**
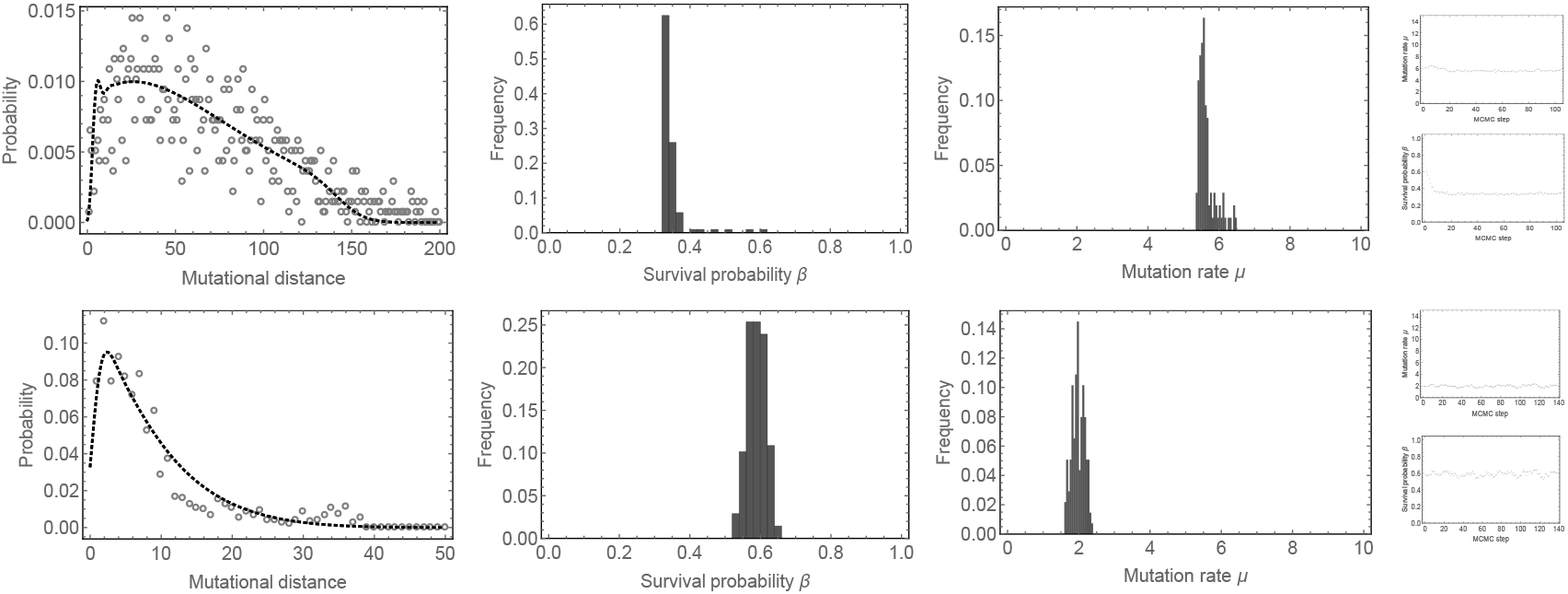
Mutational distance distribution and MCMC inference from 2 TracerX patients^28^. Note, the first case has an approximately 5 times increased distance of mutational distances compared to most colon, renal and lung cases analysed here. Together with the MSI colorectal cancer this patient has the highest mutation rate per cell division. Similar to the MSI colorectal cancer, this patient also presents with low per-cell survival probability, suggesting more cell death and cell turn over compared to other cases.

**SI Figure 19:**
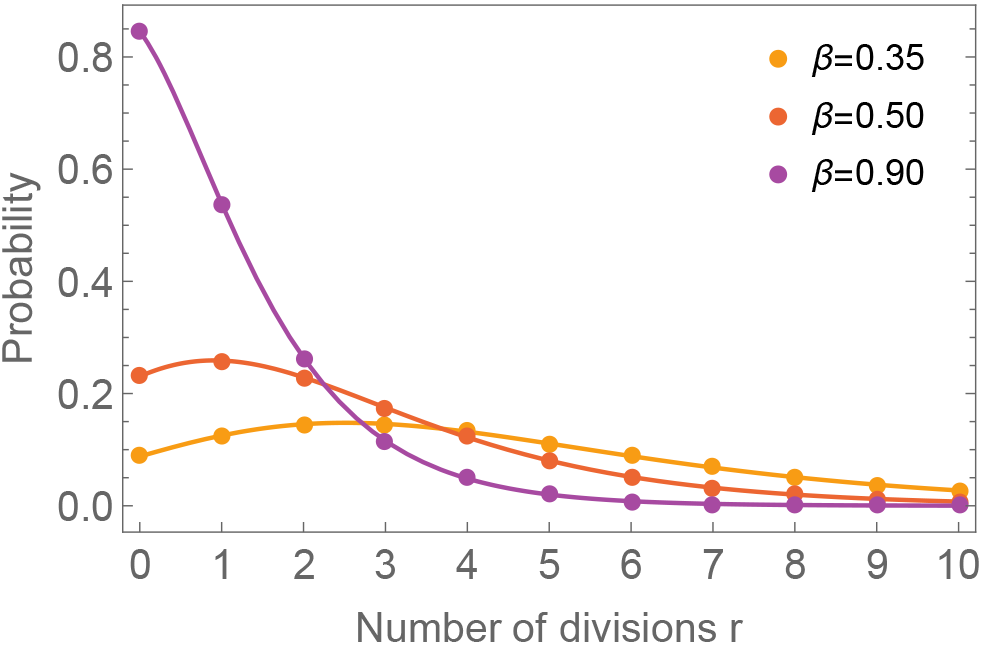
Distribution of cell divisions r in an exponentially growing population according to Equation (18).

**SI Figure 20:**
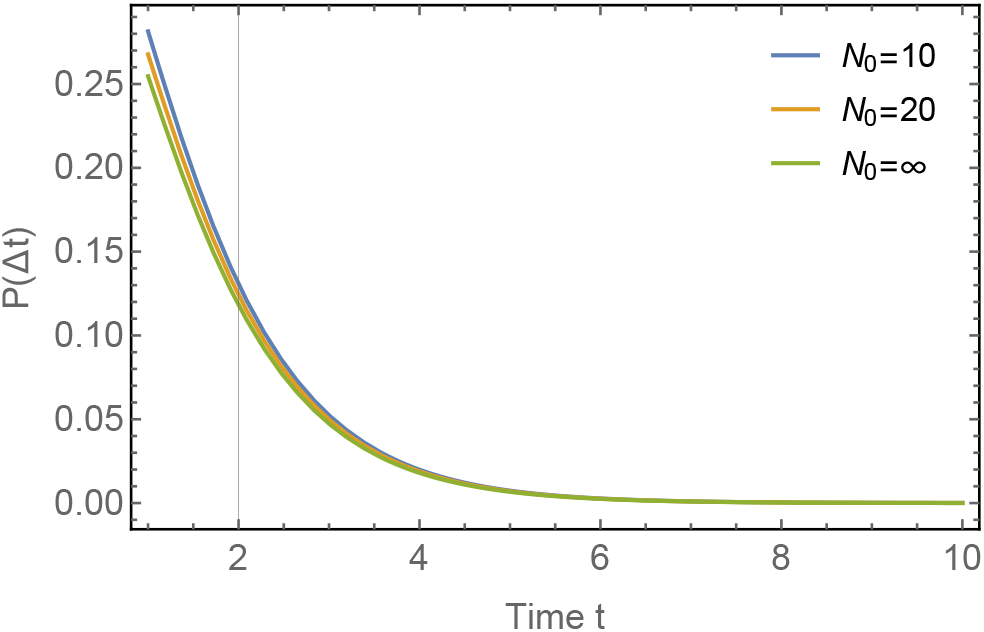
Approximation of the probability of coalescence time differences. Shown is the approximation 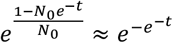 for different values of *N*_0_. The approximation works well even for small *N*_0_.

**SI Figure 21:**
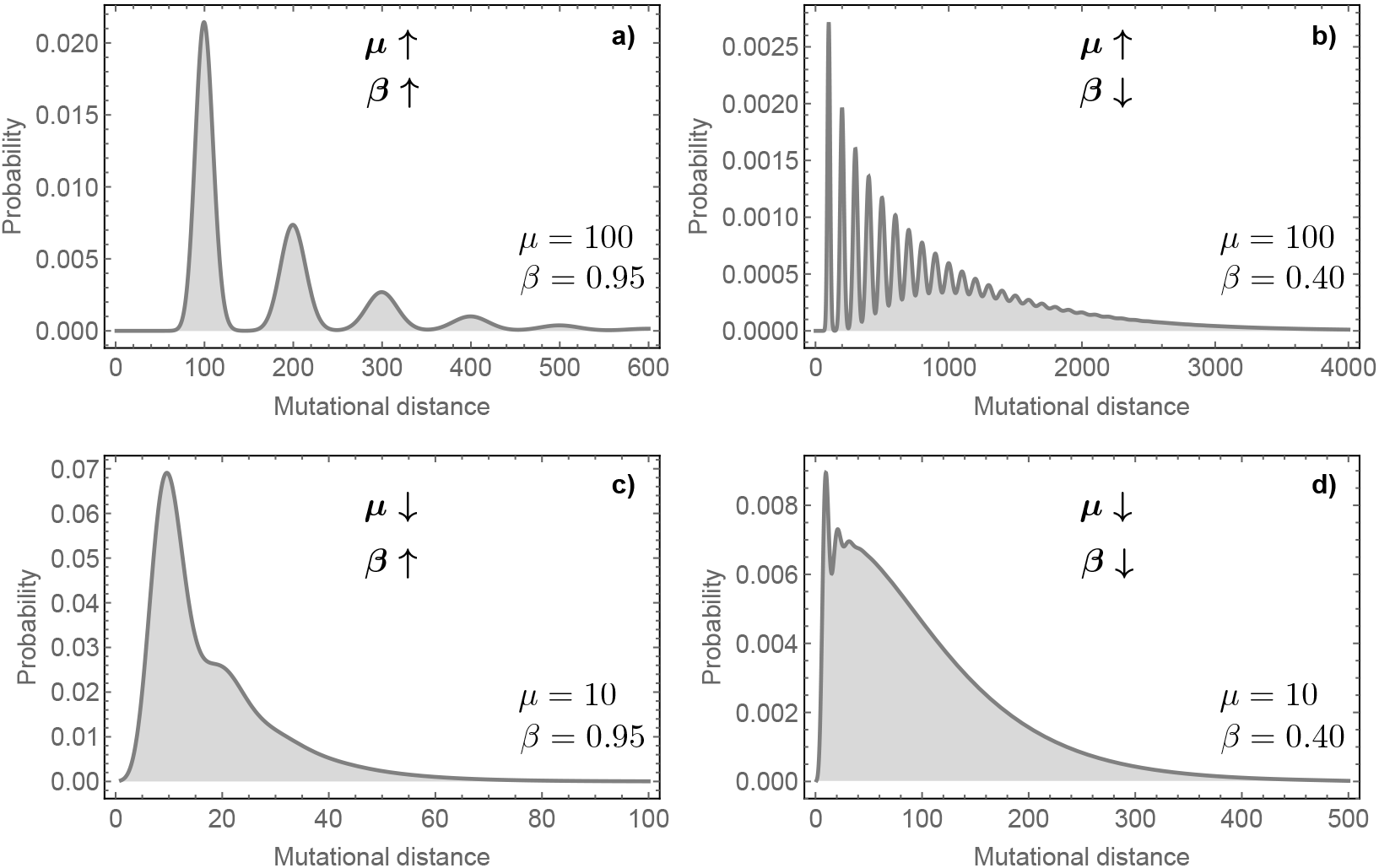
Predicted shapes of the mutational distance distribution for different combinations of small and large *μ* and *β*. The expected shape of the distribution differs greatly between different combinations of parameters. **(a,b)** Multi-modality is evident for sufficiently large mutation rates per cell division. **(c,d)** Uni-modality becomes dominant for small mutation rates per cell division. Furthermore, the length of the tail as well as the height of the distribution is largely determined by the per-cell survival rate.

**SI Figure 22:**
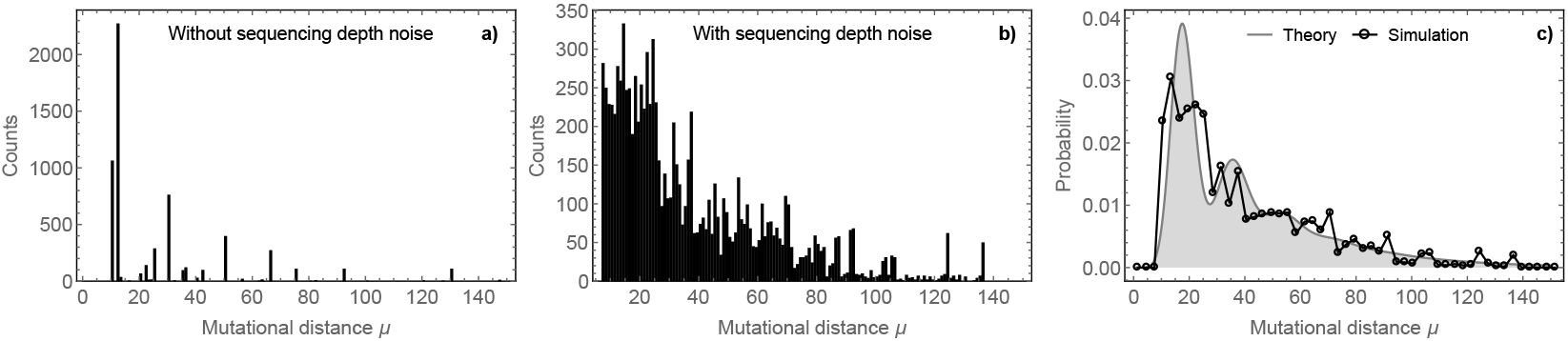
Mutational distance distribution in idealised and realistic data. **(a)** We show the mutational distance distribution inferred from a single stochastic simulation with *μ* = 15 and *β* = 0.8 for a situation with perfect clonal information, no lineage intermixing and no sequencing noise. In this situation repeated sampling of pairwise mutational distances only finds a limited number of discrete distributed peaks. Discrete peaks and a declining tail are hinted, but the expect theoretical distribution is not obvious. **(b)** Mutational distance distribution reconstructed from a simulation with same parameters, but realistic spatial sampling of intermixed lineages and sequencing depth noise. Many more unique mutational distances are evident. The distribution remains noisy (it is derived from 9 bulk samples). **(c)** The same distribution as in **(b)** (black dots), also noisy allows reconstructing the theoretically expected mutational distance distribution and to infer the underlying evolutionary parameters *μ* and *β*.

**SI Figure 23:**
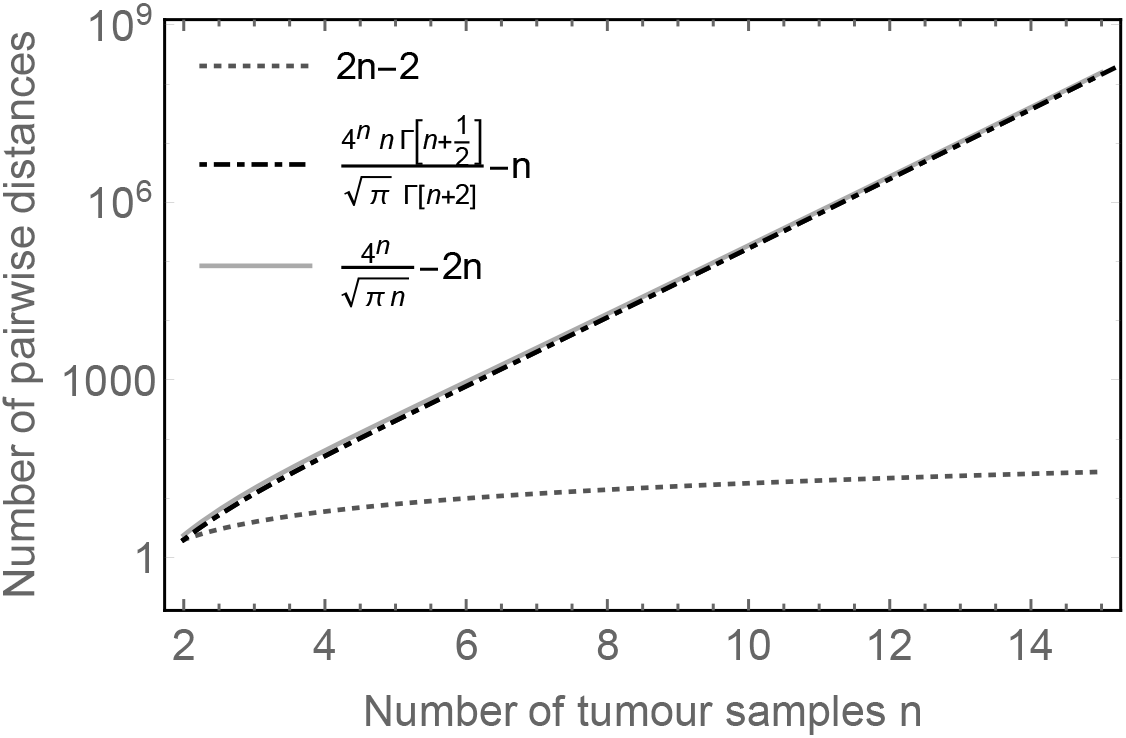
Number of sampled distances. The number of intermittent branches of a time ordered binary tree is 2*n* − 2 and scales linearly in the number of available tumour samples *n* (grey line). In contrast, the number of pairwise distances scales ~4^*n*^ and thus exponential in the number of tumour samples *n*. The exact expression is given by black dash-dotted line, whereas the light grey line shows an approximation for sufficiently large *n*. The number of pairwise differences scales very fast with the number of samples n. This is not a problem for current bulk tumour samples where *n* ≈ *7 to* 13. However, in the case of healthy haematopoiesis *n* = 89 samples are available and thus calculating all pairwise differences is impossible. However, in that case we are interested in early branching and thus restricted our analysis to distances between the first 16 identifiable ancestral cells.

**SI Figure 24:**
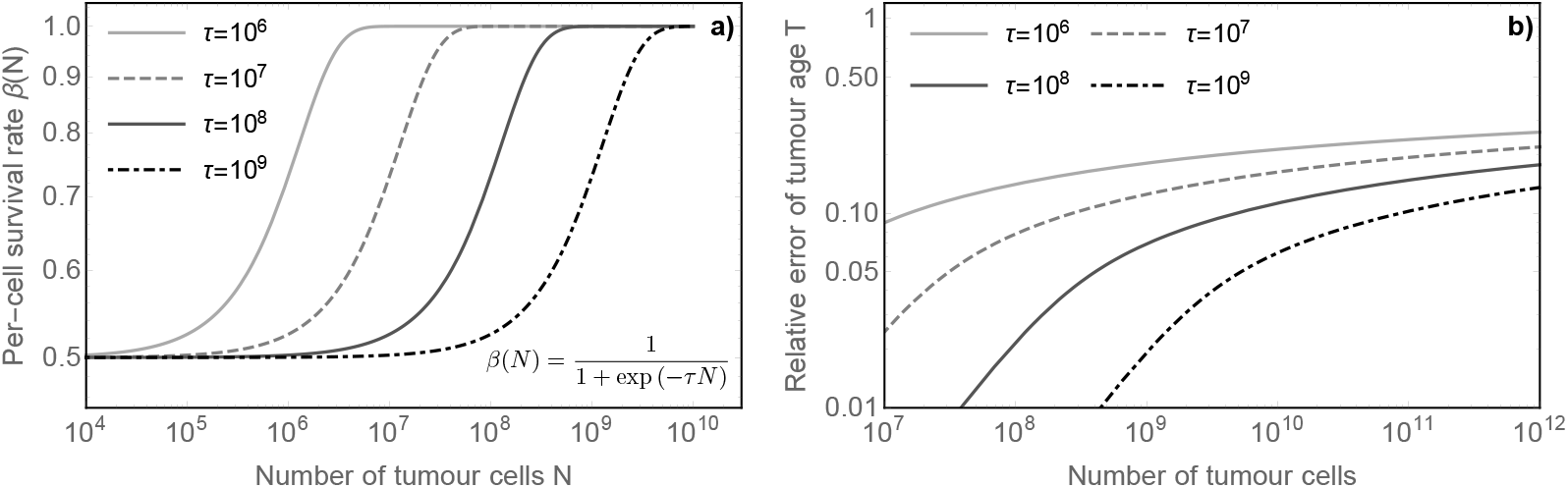
Relative error of tumour age estimates for time dependent cell-survival parameter *β*. **(a)** Shape of the Fermi-function to model the time dependence of *β* for different scale parameters *τ*. In our example, cells start with survival probability 1/2 and will acquire the maximal survival probability of 1 eventually. **(b)** Relative error of tumour age estimates, given different *τ* parameters and different times (tumour size) of diagnosis *N*_*D*_. Even if a tumour has acquired a maximal per-cell survival probability of 1 already when only 1 million tumour cells are present (the current detection threshold is approx. 100 million cancer cells), the relative error remains <20%.

**SI Figure 25:**
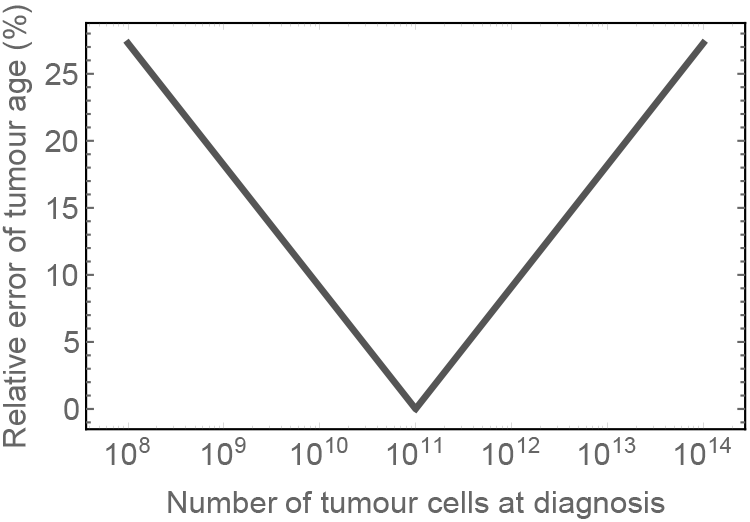
Relative error of tumour age inferences based on the tumour size at diagnosis. Plotted is 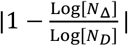 over *N*_Δ_, the actual tumour age at diagnosis, if for the calculation of tumour age a size of *N*_*D*_ = 10^11^ cells was assumed. Deviations of one order of magnitude of tumour size at diagnosis correspond to approx. 10% error for the estimation of tumour ages.

**SI Figure 26:**
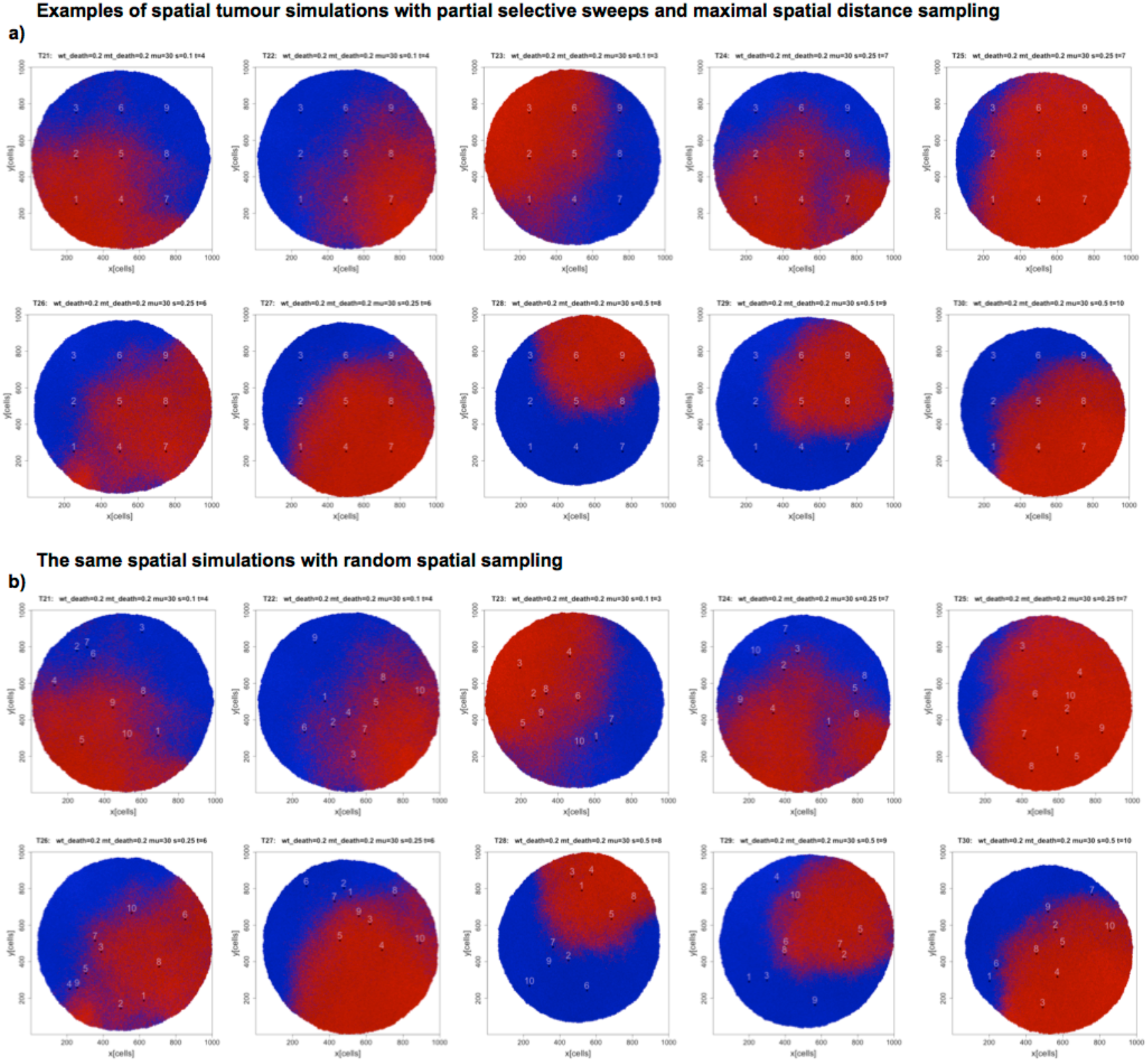
Examples of spatial simulations with partial selective sweeps and varying selection strength (s=0.1 to s=0.5). Positively selected cells are shown in red, background wild type cells in blue. Each cell carries up to thousands of private mutations accumulated during stochastic growth that are not indicated by colour here. Small areas and numbers correspond to the locations of bulk samples. **(a)** Maximal distance sampling strategy. **(b)** Randomly placed sampling strategy. In each case 9 samples were used for the construction of the mutational distance distribution and the parameter inference.

**SI Figure 27:**
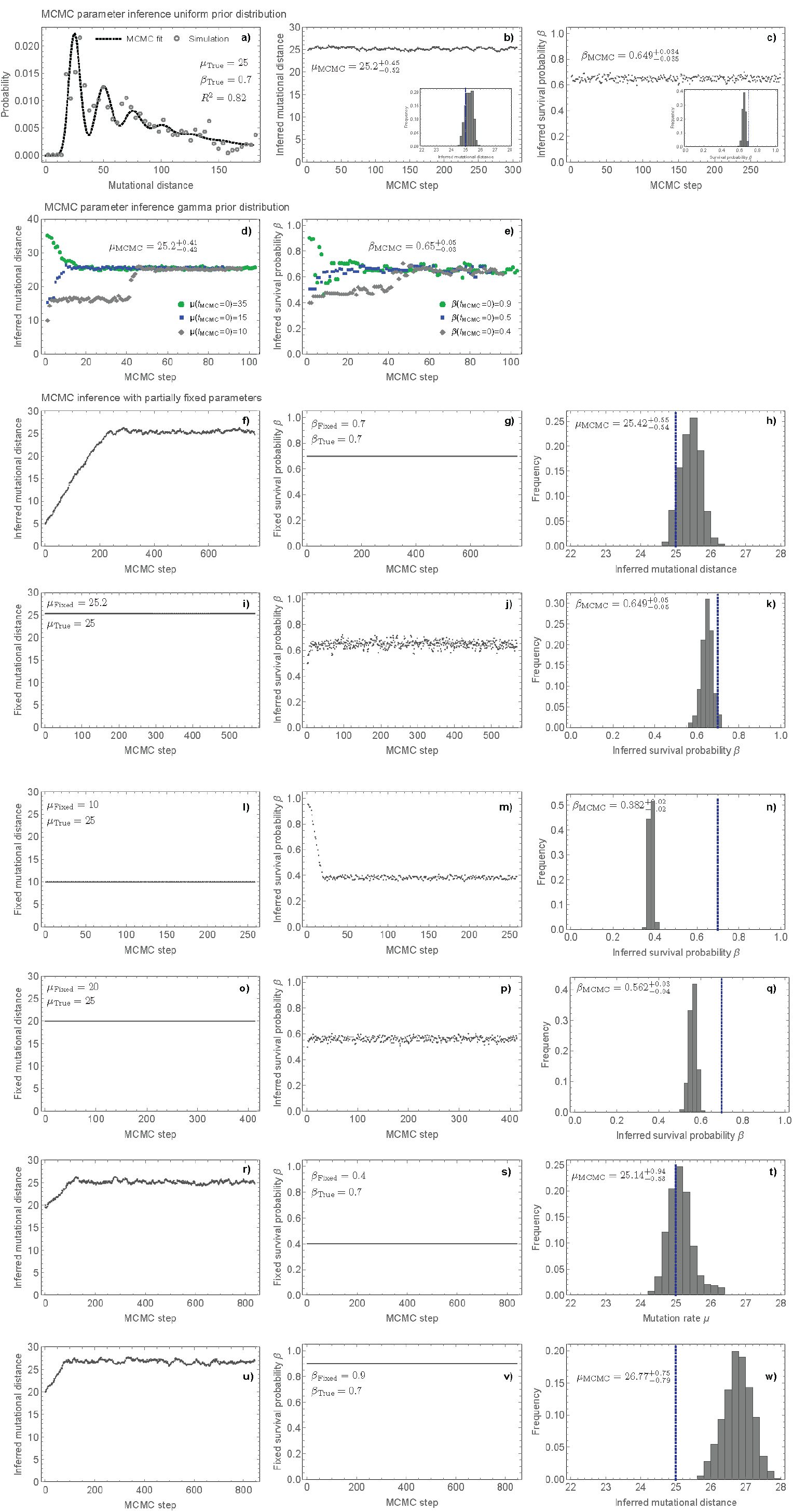
MCMC parameter inference. **(a)** Shown is one example of the mutational distance distribution inferred from 9 bulk samples of one stochastic spatial simulation of tumour growth. Subsequent panels show different scenarios for the MCMC inference, based on uniform prior **(b,c)**, Gamma prior **(d,e)** and different scenarios where one of the parameters is fixed a priori **(f-w)**.

**SI Figure 28:**
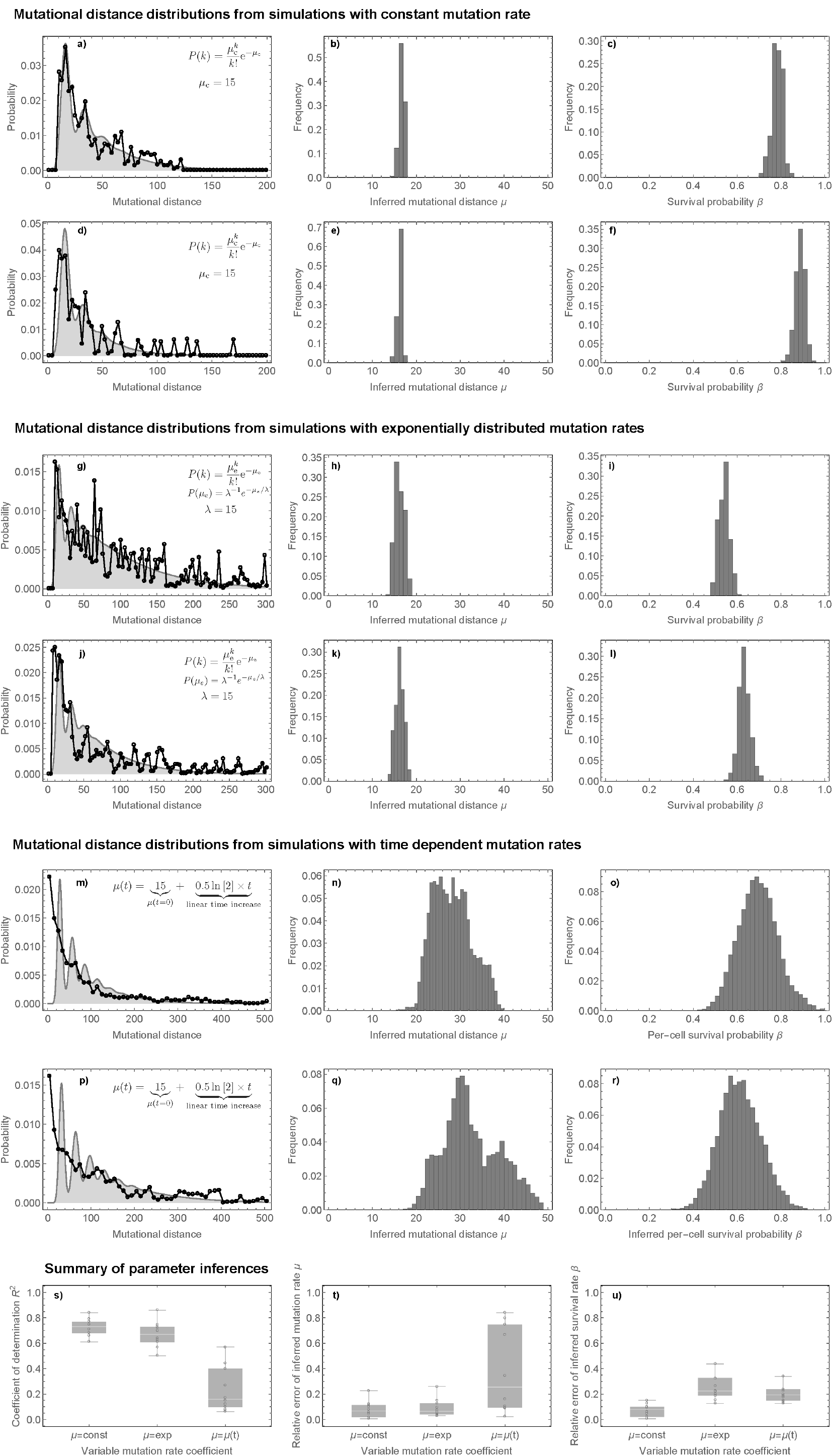
Robustness of the method of mutational distances to changes in mutation rate over time. Examples of mutational distance distributions and corresponding parameter inferences reconstructed from 9 bulk samples of stochastic spatial simulations of tumour growth with different modes of mutation accumulation. **(a-f)** Mutational distances and parameter inferences for two representative cases of simulated tumours with standard Poisson mutation accumulation. **(g-l)** Mutational distances and parameter inferences for two simulated tumours where the mutation rate is itself a random variable (here with an exponential distribution). **(m-r)** Mutational distances and parameter inferences for simulated tumours where the mutation rate increases linearly in time. In all these cases the ground truth mutation rate was 15 and survival rate was 0.8. Goodness of fit measure **(s)**, absolute error in the estimation of the per-cell mutation rate **(t)** and the per-cell survival rate **(u)** for the three different models (20 simulated tumour instances per scenario).

**SI Figure 29:**
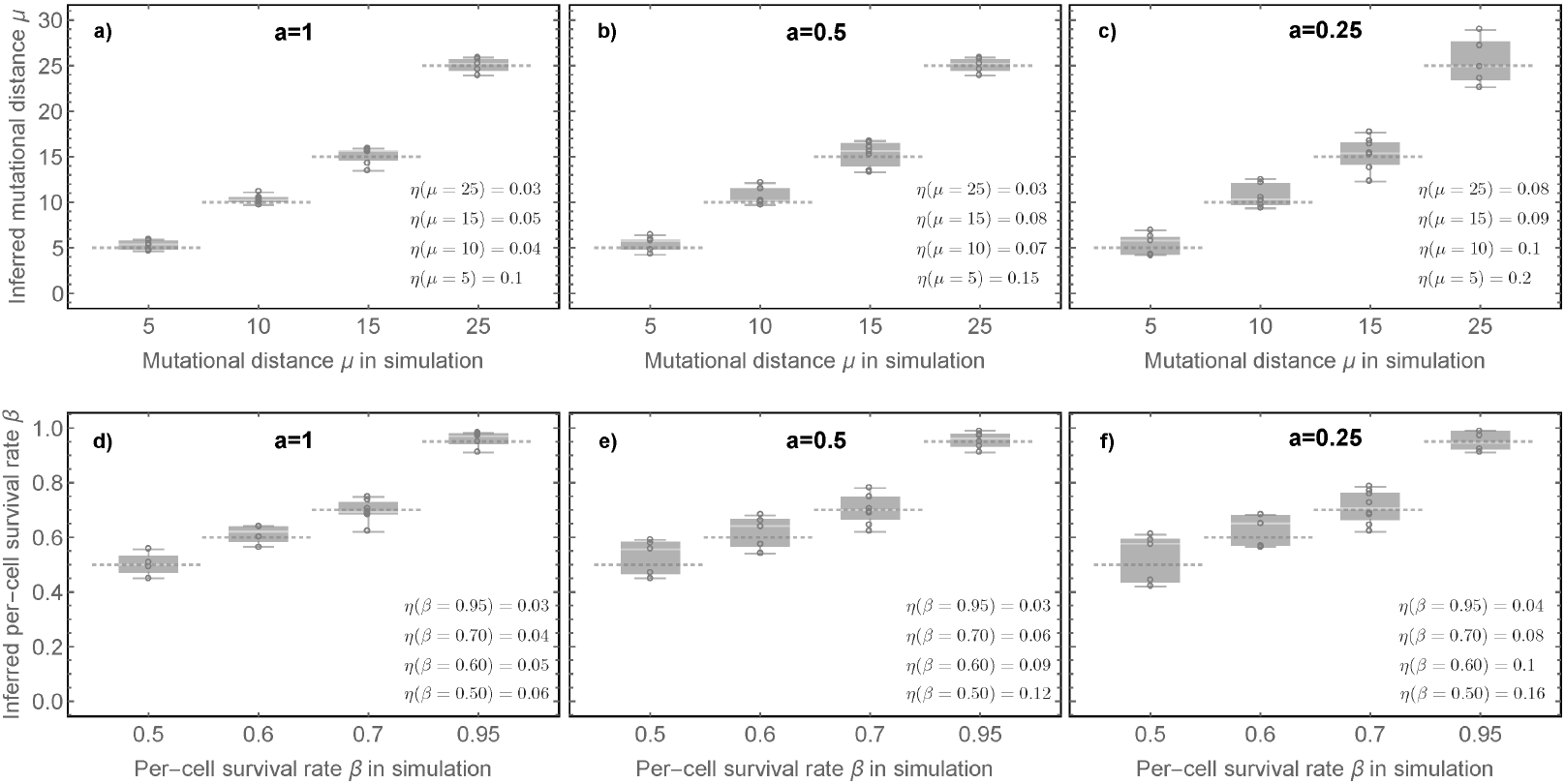
Robustness to non-exponential tumour growth. **(a)** If all cells are allowed to proliferate within a tumour, this gives rise to exponential growth. **(b)** Peripheral or ‘boundary driven’ growth leads to polynomial expansion instead. **(c-e)** Mutational distance inferences for the aggression coefficient *a* = 1, *a* = 0.5 and *a* = 0.25 (probability to proliferate in the absence of empty space). **(f-h)** Per-cell survival rate inferences. Dashed lines show ground truth, dots represent parameter inference from one stochastic spatial simulation and 9 independent bulk samples. The relative error *η* is shown in each panel.

**SI Figure 30:**
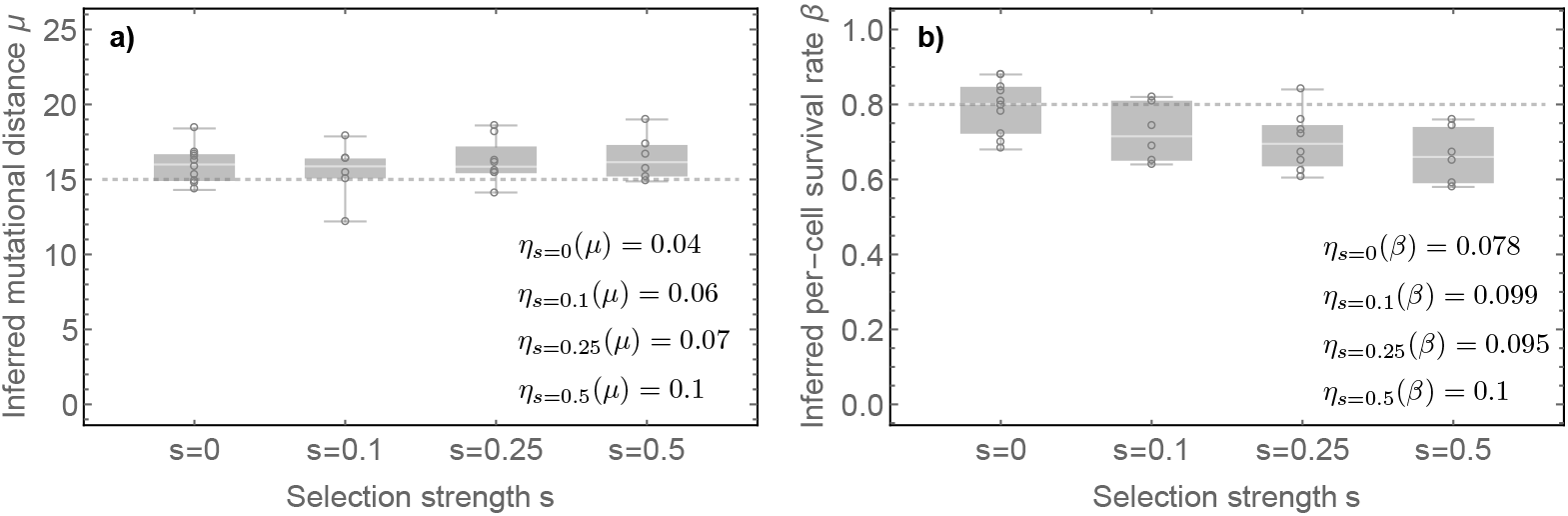
Robustness to partial selective sweeps. Parameter estimates of the **(a)** per-cell mutation rate and **(b)** per cell survival rate for a series of stochastic spatial simulations of tumour growth with partial selective sweeps and varying selection strength *s*. In our simulations *s* = 0 corresponds to the absence of positively selected clones. A clone with *s* = 0.5 proliferates 50% faster compared to background clones and thus rises in frequency over time. Examples of these simulated tumours are shown in SI Figure 24. For each parameter estimate 9 bulk samples were used to reconstruct the mutational distance distribution. Dashed lines show the exact parameters imposed on the simulations. The relative error *η* is shown in both panels.

**SI Figure 31:**
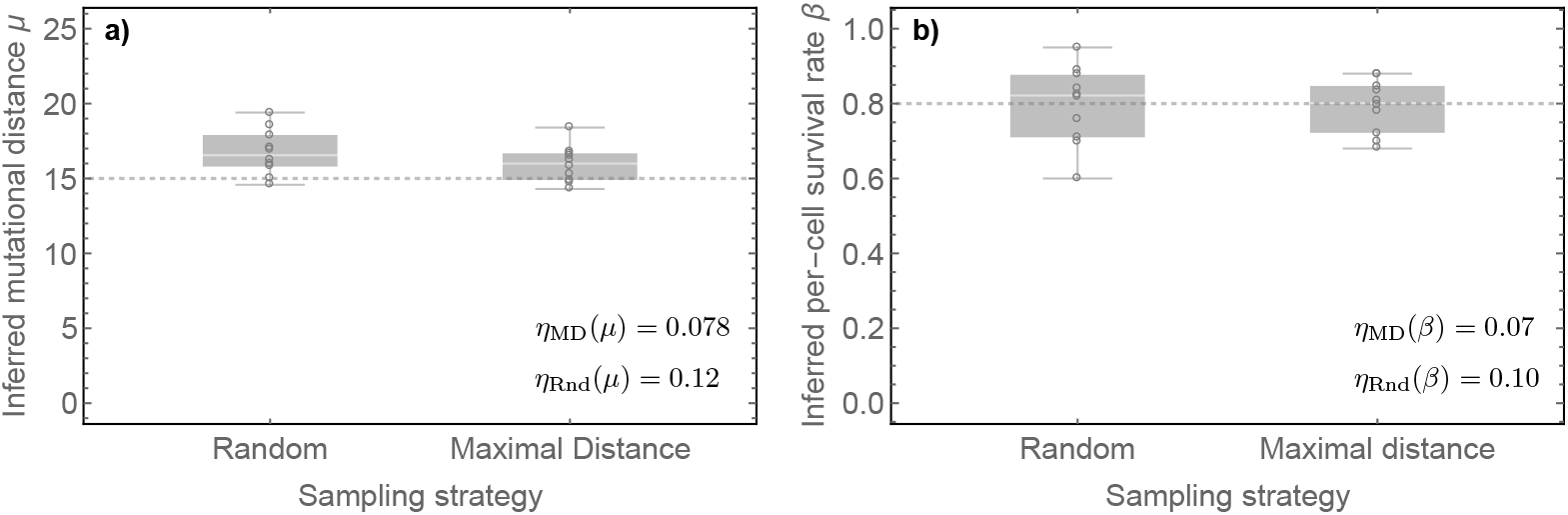
Robustness to different spatial sampling strategies. For parameter inferences 9 bulk samples per tumour were used to construct the mutational distance distribution. Examples for the different sampling strategies are shown in SI Figure 24. **(a)** absolute and relative errors in the estimation of the per-cell mutation rate **(a)** and per-cell survival rate **(b)** for random vs maximal distance sampling strategies.

**SI Figure 32:**
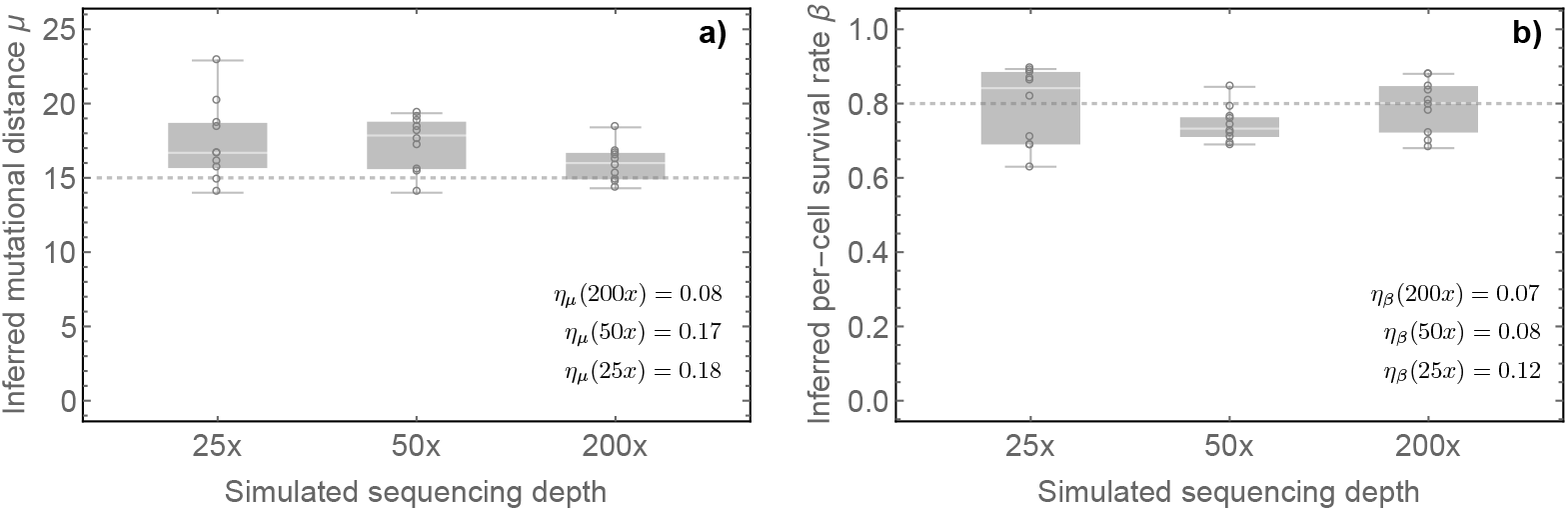
Parameter inference is robust to sequencing depth. Shown are the parameter inferences of the mutation rate **(a)** and the survival rate **(b)** for 10 spatial tumour simulations with *μ* = 15 and *β* = 0.8 from the mutational distance distribution derived from 9 bulk samples with simulated sequencing depth of 200x, 50x and 25x. Shown are also the relative errors *η* for each scenario.

**SI Figure 33:**
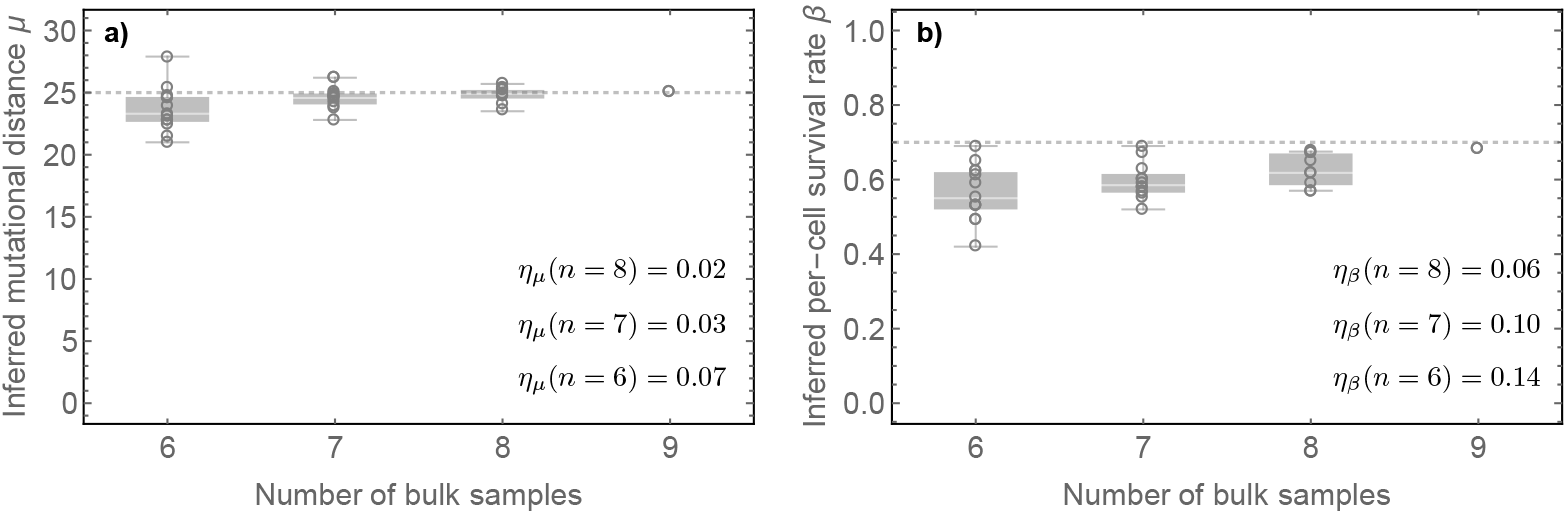
Robustness of parameter inference for decreasing number of bulk samples. Shown are inference results for stochastic spatial tumour simulations and a different number of bulk samples analysed. Decreasing the number of bulk samples increases parameter uncertainties, but the inferences remain robust for up to 6 bulk samples.

**SI Figure 34:**
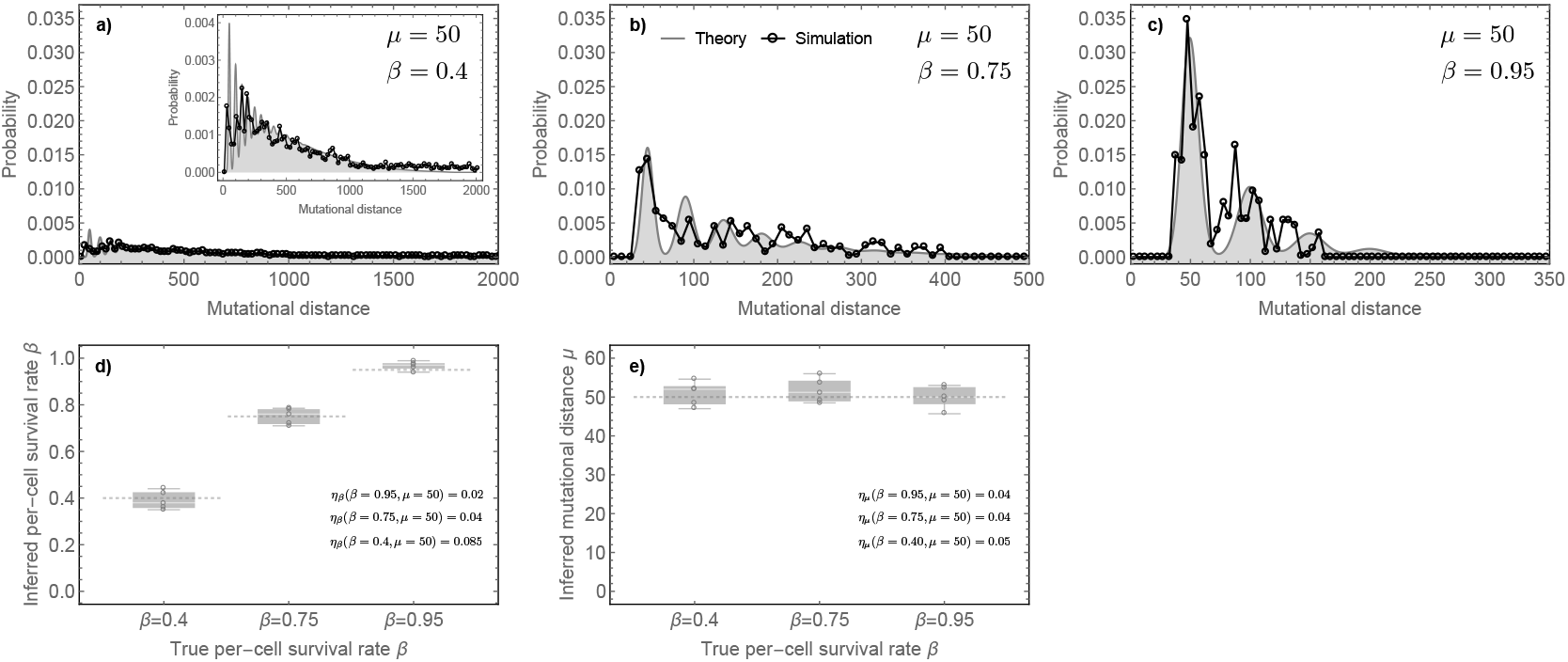
Spatial stochastic simulations with high mutation rate per cell division and different per-cell survival rates. Panels **(a)-(c)** show examples for the mutational distance distribution reconstructed for cases of high mutation rate and different per-cell survival rates. The distributions are plotted with same y-axes to show the dramatic differences in the shape of the distributions (notice the different scales of x-axis thought). The inset of panel **(a)** shows the same distribution, just with a differently scaled y-axis. Panels **(d) & (e)** show the inference of the evolutionary parameters for independent stochastic runs of spatial tumour simulations (9 bulk samples per simulation). Inferences are robust for low and high death and high mutation rates as shown by the small relative errors *η*.

**SI Figure 35:**
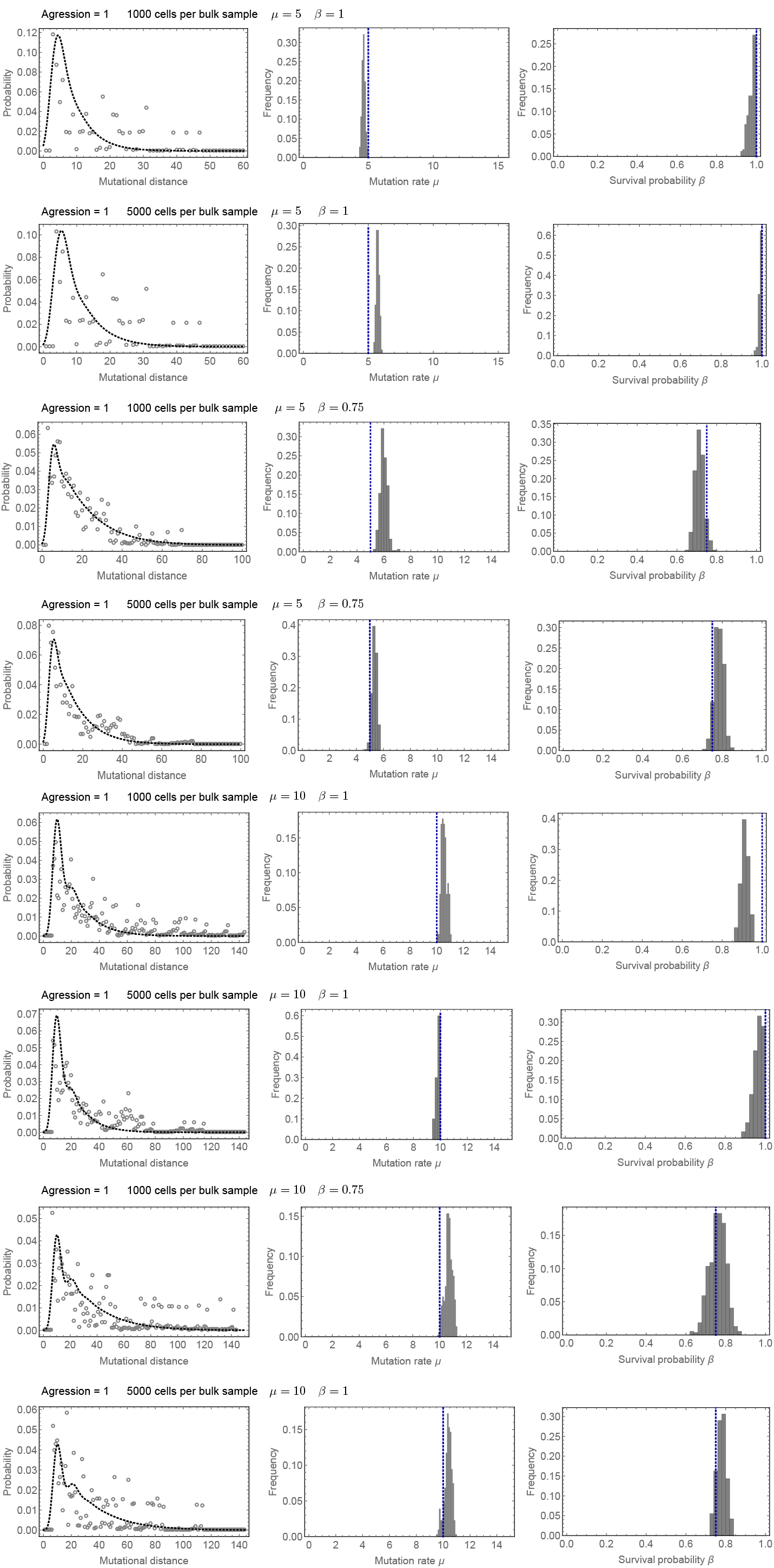
Mutational distance distribution and MCMC inference from stochastic individual based simulated tumours. Dashed lines show true parameter values. Parameter inference clusters around true values, see also Figure 1 in the main text for a summary of all inferred parameter values.

**SI Figure 36:**
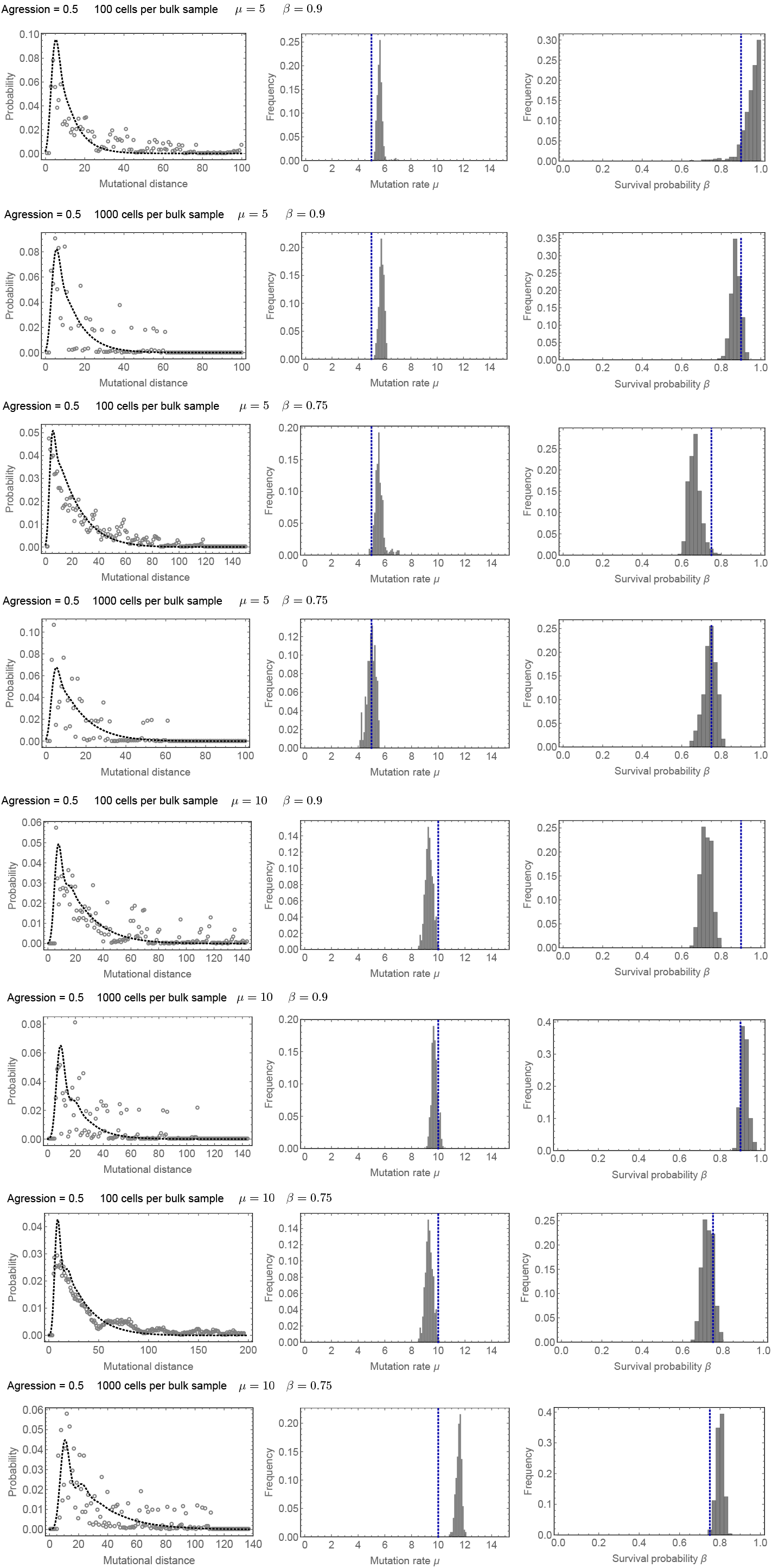
Mutational distance distribution and MCMC inference from a second set of stochastic individual based simulated tumours. Dashed lines show true parameter values. Parameter inference clusters around true values, see also Figure 1 in the main text for a summary of all inferred parameter values.

## References

1. Greaves, M. & Maley, C. C. Clonal evolution in cancer. Nature 481, 306–313 (2012).

2. Baca, S. C. et al. Punctuated Evolution of Prostate Cancer Genomes. Cell 153, 666–677 (2013).

3. Swanton, C. Intratumor Heterogeneity: Evolution through Space and Time. Cancer Research 72, 4875–4882 (2012).

4. Marusyk, A. et al. Non-cell-autonomous driving of tumour growth supports sub-clonal heterogeneity. Nature 514, 54–58 (2014).

5. Lengauer, C., Kinzler, K. W. & Vogelstein, B. Genetic instability in colorectal cancers. Nature 386, 623–627 (1997).

6. McGranahan, N. & Swanton, C. Biological and Therapeutic Impact of Intratumor Heterogeneity in Cancer Evolution. Cancer Cell 27, 15–26 (2015).

7. McGranahan, N. & Swanton, C. Clonal Heterogeneity and Tumor Evolution: Past, Present, and the Future. Cell 168, 613–628 (2017).

8. Burrell, R. A., McGranahan, N., Bartek, J. & Swanton, C. The causes and consequences of genetic heterogeneity in cancer evolution. Nature 501, 338–345 (2013).

9. Lynch, M. Evolution of the mutation rate. Trends in Genetics 26, 345–352 (2010).

10. Lynch, M. et al. Genetic drift, selection and the evolution of the mutation rate. Nature Reviews Genetics 17, 704–714 (2016).

11. Chen, X. et al. Single-cell analysis at the threshold. Nature Biotechnology 34, 1111–1118 (2016).

12. Kuipers, J., Jahn, K. & Beerenwinkel, N. Advances in understanding tumour evolution through single-cell sequencing. Biochimica et Biophysica Acta 1867, 127–138 (2017).

13. Davis, A. & Navin, N. E. Computing tumor trees from single cells. Genome Biology 1–4 (2016). doi:10.1186/s13059-016-0987-z

14. Sottoriva, A. et al. A Big Bang model of human colorectal tumor growth. Nature Genetics 47, 209–216 (2015).

15. Williams, M. J., Werner, B., Barnes, C. P., Graham, T. A. & Sottoriva, A. Identification of neutral tumor evolution across cancer types. Nature Genetics 48, 238–244 (2016).

16. Graham, T. A. & Sottoriva, A. Measuring cancer evolution from the genome. Journal of Clinical Investigation 241, 183–191 (2016).

17. Williams, M. J. et al. Quantification of subclonal selection in cancer from bulk sequencing data. Nature Genetics 1–14 (2018). doi:10.1038/s41588-018-0128-6

18. Bozica, I. et al. Accumulation of driver and passenger mutations during tumor progression. Proceedings of the National Academy of Science 107, 18545–18550 (2010).

19. Milholland, B. et al. Differences between germline and somatic mutation rates in humans and mice. Nature Communications 8, 1–8 (2017).

20. Lee-Six, H. et al. Population dynamics of normal human blood inferred from somatic mutations. Nature 1–18 (2018). doi:10.1038/s41586-018-0497-0

21. Cross, W. et al. The evolutionary landscape of colorectal tumorigenesis. Nature Ecology & Evolution 1–14 (2018). doi:10.1038/s41559-018-0642-z

22. Frigola, J. et al. Reduced mutation rate in exons due to differential mismatch repair. Nature Genetics 1–13 (2017). doi:10.1038/ng.3991

23. Brody, Y. et al. Quantification of somatic mutation flow across individual cell division events by lineage sequencing. Genome Res. 28, 1901–1918 (2018).

24. Alexandrov, L. B. et al. Signatures of mutational processes in human cancer. Nature 500, 415–421 (2013).

25. Alexandrov, L. B. et al. Clock-like mutational processes in human somatic cells. Nature Genetics 47, 1402–1407 (2015).

26. Ebersberger, I., Metzler, D., Schwarz, C. & Pääbo, S. Genomewide Comparison of DNA Sequences between Humans and Chimpanzees. Am. J. Hum. Genet. 70, 1490–1497 (2002).

27. Roerink, S. F. et al. Intra-tumour diversification in colorectal cancer at the single-cell level. Nature 1–22 (2018). doi:10.1038/s41586-018-0024-3

28. Jamal-Hanjani, M. et al. Tracking the Evolution of Non–Small-Cell Lung Cancer. New England Journal of Medicine 376, 2109–2121 (2017).

29. Gerlinger, M. et al. Genomic architecture and evolution of clear cell renal cell carcinomas defined by multiregion sequencing. Nature Genetics 46, 225–233 (2014).

30. Benatti, P. Microsatellite Instability and Colorectal Cancer Prognosis. Clinical Cancer Research 11, 8332–8340 (2005).

31. Vilar, E. & Gruber, S. B. Microsatellite instability in colorectal cancer—the stable evidence. Nat Rev Clin Oncol 7, 153–162 (2010).

32. Hause, R. J., Pritchard, C. C., Shendure, J. & Salipante, S. J. Classification and characterization of microsatellite instability across 18 cancer types. Nature Publishing Group 1–11 (2016). doi:10.1038/nm.4191

33. Slatkin, M. & Hudson, R. R. Pairwise Comparisons of Mitochondrial DNA Sequences in Stable and Exponentially Growing Population. Genetics 129, 555–562 (1991).

34. Waclaw, B. et al. A spatial model predicts that dispersal and cell turnover limit intratumour heterogeneity. Nature (2015). doi:10.1038/nature14971

35. Temko, D., Tomlinson, I. P. M., Severini, S., Schuster-Böckler, B. & Graham, T. A. The effects of mutational processes and selection on driver mutations across cancer types. Nature Communications 1–10 (2018). doi:10.1038/s41467-018-04208-6

36. Gehring, J. S., Fischer, B., Lawrence, M. & Huber, W. SomaticSignatures: inferring mutational signatures from single-nucleotide variants. Bioinformatics 31, 3673–3675 (2015).

37. Bozic, I. et al. Evolutionary dynamics of cancer in response to targeted combination therapy. eLife 2, e00747 (2013).

38. Werner, B. et al. The Cancer Stem Cell Fraction in Hierarchically Organized Tumors Can Be Estimated Using Mathematical Modeling and Patient-Specific Treatment Trajectories. Cancer Research 76, 1705–1713 (2016).

39. Werner, B., Traulsen, A., Sottoriva, A. & Dingli, D. Detecting truly clonal alterations from multi-region profiling of tumours. Scientific Reports 7, 1–9 (2017).

